# A Framework for Transparent and Repeatable Species Range Maps

**DOI:** 10.1101/2025.05.14.653500

**Authors:** Nathan M. Tarr

## Abstract

Knowledge of species’ geographic distributions is essential to wildlife conservation and management. A great deal of research has developed predictive models that reveal complex spatial patterns and temporal dynamics of species’ occurrence probabilities. However, binary maps of the geographic limits of species occurrence (“range maps”) are still used in conservation assessments, decision making, and as a component of the species distribution modelling process itself. Organizations that create and revise range maps face numerous challenges. For example, different types of data must often be combined to fill knowledge gaps. Range delineation is labor intensive and subjective in ways that hinder transparency and repeatability. I developed a framework for transparent and repeatable range maps in support of the U.S. Geological Survey’s National Gap Analysis Project. The framework has an explicit conceptual foundation that supports a compilation process built with open-source technologies and freely available data. As a case study, I mapped Fisher *Pekania pennanti* range and presence between 2021 and 2025 for the conterminous United States by compiling information from a range map circa 2001, 2,287 species occurrence records from 16 datasets, and 17 scientific publications. The maps depict presence and range according to levels of evidence. The justification for classification of each spatial unit is included in the output. The maps revealed that translocations and emigration have expanded the species’ range in some areas, but the species is suspected to be absent from some areas previously considered range. The framework overcomes several challenges, but some components of the process warrant further development, such as occurrence record evaluation and direct incorporation of predictions from species distribution models. This work demonstrates how freely accessible technologies and data can support the maintenance and revision of species range datasets while improving transparency, reproducibility, and efficiency.

## Introduction

Knowledge of species’ geographic distributions is essential to conservation efforts. Animal populations, the resources they depend upon, and the threats they face are distributed unevenly across space, thus it is important to assess the spatial overlap of species and threats. Spatial ecologists often seek analytical methods for mapping species distributions that reveal and reflect ecological complexity because the processes and relationships that underly species’ distributions are complex and uncertain. However, maps defining geographic limits remain valuable for researchers and decision makers who apply spatial data (Brown et al. 1996; Jetz et al. 2012; Jetz et al. 2019). One example is the coarse, binary map of the limits of a species geographic distributions that are applied in conservation assessments, which are often referred to as “range maps”.

Range maps have several applications. They provide general descriptions of species’ biogeography in field guides and species accounts (NatureServe 2023; USDA, NRCS 2023); introductory content of presentations and scientific publications; and species conservation assessments (Graham and Hijmans 2006; Jetz et al. 2008). They can also provide a basis for biodiversity and conservation studies (Di Marco et al. 2017). For example, Jenkins et al. (2015) and Marsh et al. (2022) created maps of species richness from the range maps of hundreds of species. Davidson et al. (2021) incorporated range area as a metric of species rarity in an assessment of species conservation. Furthermore, the IUCN Red List uses the area of species ranges in their conservation status assessments (IUCN Standards and Petitions Committee 2022), and range maps are integral to the U.S. Fish and Wildlife Service Species Status Assessment Framework (USFWS 2016; Dunn et al. 2024).

A less-obvious application of range maps is as inputs to species distribution models. They can be used to define or inform the selection of spatial extents for models with fine-to moderate-spatial resolutions (Jetz et al. 2012; Guisan et al. 2017; Barry et al. 2021; Franklin 2023). For example, the USGS Gap Analysis Project built habitat distribution models and limited output to areas within range map boundaries (McKerrow et al. 2018; Gergely et al. 2019). Similarly, Hamilton et al. (2022) built random forest distribution models for species, and constrained model outputs to areas within range delineated by various sources including BirdLife International and Handbook of the Birds of the World (2018) and IUCN (2016). Range maps may also be used to direct sampling effort in support of occupancy or abundance models with coarse spatial resolutions. For example, Krohner et al. (2022) used occupancy models and other sources of information to define a geographical extent (sampling frame) for *Pekania pennanti* (Erxleben, 1776) (Fisher) monitoring in the northern Rocky Mountains.

Range maps are created via the synthesis of information and beliefs about species’ distributions by individuals or groups of scientists (Marsh et al. 2022; Rego et al. 2024). During map compilation, data, knowledge, and hypotheses about the locations of individuals are synthesized into approximations of the geographic limits of species occurrence. Various methods can be used to create maps (see Text S1 for further discussion).

Map compilations via data synthesis often involve subjective decisions. Thus, the bases and meanings of maps may not be clear to users without clear definitions and extensive documentation. Transparency also allows users to assess the suitability of maps for their applications and to determine why one map version differs from another. Furthermore, transparency can reveal important uncertainties that arose during map compilation. Unfortunately, providing sufficient documentation for transparency is difficult because of the volume of knowledge and data that must be synthesized, as well as the number of decisions involved in their synthesis for the large spatial, and sometimes temporal, extents that are characteristic of range maps (Jetz et al. 2019).

In addition to being important for map users, transparency is important for producers of range maps involved in long-term efforts to revise and maintain data. Map producers need the ability to explain the maps to users and reviewers during peer-review. Additionally, they must also be able to understand the provenance and basis of the maps they must revise to collaborate effectively. Without documentation, maps must be created from scratch at every iteration with limited ability to ensure that accuracy is not reduced, important past changes are not overridden, or that valuable data and knowledge are not overlooked and discarded with each revision.

Range dynamics present another core challenge to the revision of range maps over time. Climatic changes, species interactions, human activities, and landscape changes may drive local changes in occupancy that scale up to range shifts, contractions, or expansions (Jones et al. 2023). However, once areas are classified as part of a species’ range, it is difficult to gain sufficient evidence of absence to justify their removal (Jones et al. 2023). This is because range map scales are coarse and, thus, spatial units are large. Spatially and temporally complete sampling within a spatiotemporal unit is necessary to directly establish absence, extirpation, or retractions (Jetz et al. 2019). This challenge, coupled with the relative ease of adding areas to a species’ range based on few data, increases the risk of erroneous depictions of range expansions when the species’ distribution has merely shifted. Areas of range can be removed based on comprehensive evaluation of information, including expert opinion and model predictions, but as stated above, it is important to document the justification for such decisions to support users and producers of maps.

My objective was to develop a framework for long-term maintenance and revisions of species range data for the U.S. Geological Survey’s Gap Analysis Project (GAP). The Gap Analysis Project uses 30 x 30 m resolution habitat maps to estimate the proportion of habitat in different protection categories for each terrestrial vertebrate species occurring within the United States and to map patterns of species richness at a moderate spatial resolution. The recent modeling extent has been the conterminous United States (CONUS). The habitat maps are constrained to areas within species’ range limits. I sought to design a range-mapping framework that allows the synthesis and integration of observation data, expert opinion, information from scientific publications, and predictive models. The framework needed to provide sufficient documentation for transparency and facilitate repeated compilation to support iterative refinement of maps in the short term, as well as revisions and updates over time. I also sought to create an instance of the framework with open-source software.

## Methods

This framework generates species range and presence maps across CONUS. Maps are created on a natural grid by integrating data sources, including GAP version 1 national range data (USGS 2018), species occurrence records, and expert opinions at each spatial unit. The framework is structured with 5-year steps whereby the use of the most current and reliable data is prioritized for range delineation at each step while incorporating older data to address knowledge gaps. Two core principles guide map compilation: (1) spatial units lacking evidence of species presence are assumed to be outside the species’ range, and (2) continued species presence or range within a spatial unit is assumed unless contradicted by available evidence. Data integration follows a defined information hierarchy, incorporating weighted scores, ranks, and user-defined parameters that are explicitly documented. This process facilitates efficient and transparent map updates as new information becomes available.

### Conceptual Basis and Definitions

A universally accepted definition for range maps does not exist to distinguish them from other types of species distribution maps, especially those with coarse spatial resolutions. However, the implementation of a framework requires explicit definitions for range and associated concepts because formalized processes for map compilation must follow standardized rules.

In this framework, range is defined as, “the areas containing the vast majority of individuals of a taxon within a specified time period as represented at a coarse spatial resolution.” Presence refers to whether there is sufficient evidence that an individual of the species occurred within a period. Because range is conceptually defined by the locations of most individuals of a taxon, it is tied to the concept of geographic outliers: those which occur outside the range limits. Such extralimital individuals are considered exceptions and sometimes referred to as “vagrants”, “transients”, “rarities”, “casuals”, or “accidentals”, which are loosely defined synonyms for outliers (Lees and Gilroy 2021). Hereafter, I use the term “extralimitals” to refer to all such exceptions. Importantly, the conceptual boundary between range and non-range (i.e., the range limit) is the same as the conceptual boundary between extralimital and intralimital records (e.g., vagrant vs. non-vagrant). In practice, it is not always clear whether an observed individual is extralimital because processes for defining range limits are approximate and somewhat subjective. The identification of extralimital records is a difficult component of data-driven range map compilations.

Importantly, range definitions are sensitive to the chosen spatiotemporal resolution because individual animals move over time and binary range maps generalize those locations to spatiotemporal units. Maps with different spatiotemporal resolutions may present different spatial patterns even if the underlying data are the same. Scale is not consistent among the many range maps that have been produced to date, but range maps generally have large spatial extents, coarse spatial grain, and coarse or unspecified temporal scales. One exception is the collection of weekly range maps created by eBird (Fink et al. 2023).

The scale of GAP range data is 12-digit hydrologic units (USGS 2011, see description below) over five-year periods within CONUS and after 2001. The Gap Analysis Project’s choice of a spatial resolution for range maps was a tradeoff between detail and feasibility. Whereas finer resolutions support the avoidance of commission errors in habitat maps, coarser resolutions create lower demands for data quantity, storage, and processing.

### Inputs

#### Spatial Units

GAP developed a natural grid for the version 1 GAP range dataset (GAP HU’s; USGS 2018) that comprises 82,717 polygons covering CONUS and extending 5 km into coastal waters. The grid was derived from the USGS Watershed Boundary Dataset (Jones et al. 2022). To accommodate GAP range and habitat modeling needs, GAP adjusted polygon boundaries at the 12-digit hydrologic unit level as necessary (USGS 2011).

#### Species Occurrence Data

Large quantities of species occurrence records (i.e., observations) are freely available from the Global Biodiversity Information Facility (GBIF). These records can provide valuable evidence of past species distributions, informing the compilation of range maps. Our methods for acquiring and curating sets of species occurrence data are detailed below in *Curate Occurrence Records*.

#### Opinion Data

We developed a private database to store expert opinions on species presence and range. This database includes tables for opinions about summer, winter, and year-round range, as well as species presence. Each table records expert assessments on the presence or absence of individuals or range within spatial units, along with associated confidence levels and expert rankings (detailed in Table S1). The framework distinguishes between three types of expert opinion:

1. Primary Opinions: Based on personal experiences, such as field observations and research.
2. Secondary Opinions: Based on another expert’s opinions or research, such as knowledge and beliefs about a species range gleaned from scientific literature.
3. Tertiary Opinions: Based on general or comprehensive review of data inputs or preliminary map results. For example, if a staff member believes that a range boundary in an initial version was inaccurate because it was coarsely drawn and included areas that can now be removed at a finer resolution.

Although the compilation process does not treat the three types of opinions differently, recognizing them helps define “expert opinion” and clarifies the basis of and ways in which GAP staff and other experts can alter the range maps. Treating the results of literature reviews as secondary expert opinions is beneficial because it acknowledges that information is interpretated, synthesized, and scrutinized by the map producer.

#### GAP Version 1 Range Data

In 2018, GAP published version 1 of its CONUS-wide range maps for 1,560 species (USGS 2018). This dataset attributed each species to GAP HU’s where it occurred. For each species-spatial unit combination, four attributes were provided: presence, season, reproduction, and origin (USGS 2018). Presence conveyed whether the species was believed to occur within the HU (e.g., extant, possibly present, extirpated). The season attribute assigned a seasonal range status to HU’s (e.g., year-round, migratory, winter, or summer) to qualify in which seasons the species is present.

This framework utilizes different presence categories than that of version 1, and range is defined and handled differently. In this framework, a separate map is generated for each relevant season for each species. Each map incorporates categories of evidence (Table 1). While the frameworks exhibit some discrepancies, the version 1 dataset can still provide valuable information to fill gaps in occurrence and expert opinion data.

**Table 1.**
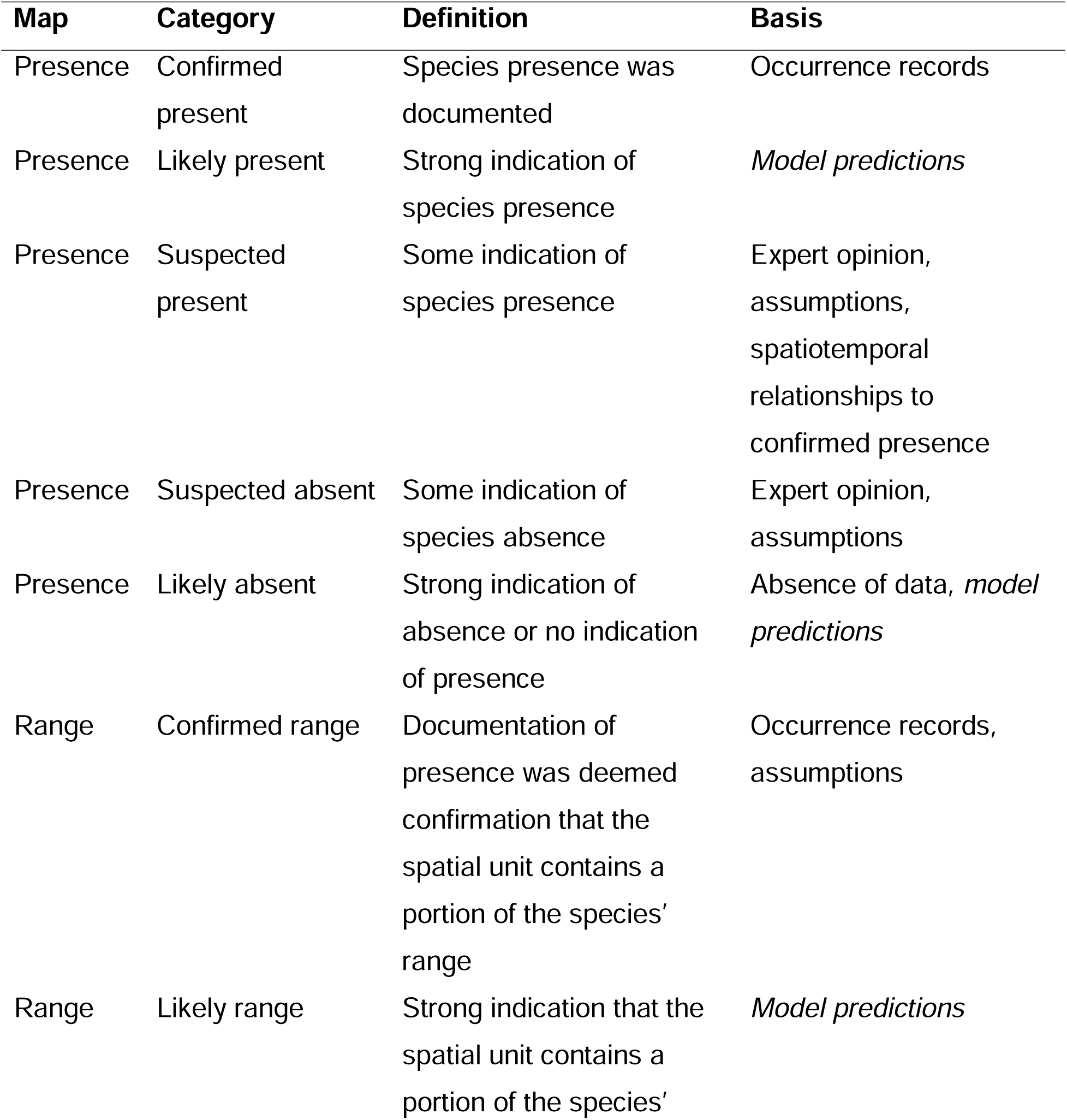
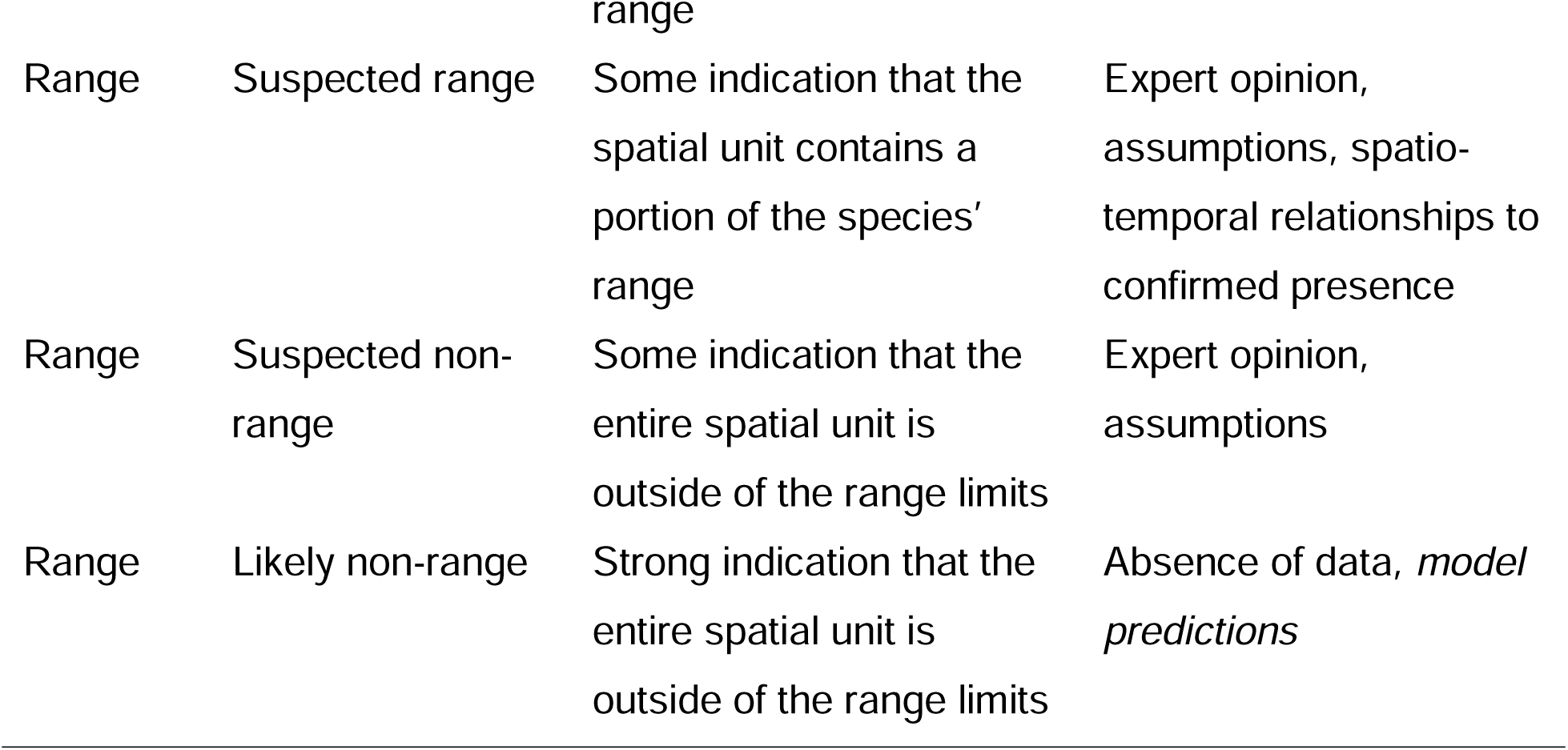
Legend categories for maps of species range and presence in the range mapping framework. A compilation process automatically assigns categories to spatial units based upon a synthesis of species occurrence records, expert opinion, and GAP version 1 range data. “Model predictions” is italicized to indicate that they are not currently incorporated into the process but will be in a future version.

#### Range Compilation Parameters

To enhance flexibility and document key decisions, we established a set of parameters to control the range compilation process. These parameters are manually entered by staff into a database at the initiation of each range compilation effort. The range compilation code retrieves the appropriate parameter set from the database during execution. The parameters encompass: 1) choices about which data to incorporate; 2) temporal filters for species observations; 3) a threshold distance beyond which records are considered extralimital; and 4) an error tolerance for assigning observations to the appropriate spatial units (Table 2).

**Table 2.** Compilation parameter list and definitions with data types for a transparent and repeatable range mapping framework. Definitions include the data type. The parameters influence the behavior of several automated subprocesses of map compilation that integrate species occurrence records, a prior range map, and expert opinions regarding species presence and range limits.

| Name | Description |
| --- | --- |
| GAP Version 1 | Whether to incorporate the GAP version 1 (2001) data.<br>Boolean. |
| Expert Opinion | Whether to incorporate opinions from the Opinion Database.<br>Boolean. |
| Observations | Whether to incorporate species occurrence records. Boolean. |
| Years | Years to include data during the compilation. List of strings. |
| Months | Months from which to include data during the compilation. List<br>of strings. |
| Extralimital<br>Distance | Threshold distance in meters from spatial units (HU's) that are<br>coded present, beyond which occurrence records are judged to<br>be extralimital. Integer. |
| Spatial Error<br>Tolerance | The maximum acceptable risk of assigning a species<br>occurrence record to a spatial unit (i.e., HU) that neighbors the<br>unit where the organism was observed. Integer. |

### Outputs

#### Range Database

The range compilation process generates a self-contained SQLite database that stores all results, as well as spatial metadata. The database comprises multiple tables (Table 3), with the primary results stored in tables named “presence,” “summer,” “winter,” and “year-round,” depending on the species’ seasonal occurrence. Presence and range are coded according to categories that reflect levels of belief that a species was present in the unit and that range occurs within the spatial unit, respectively (Table 1). Some non-range spatial units can be assigned a “present” code because of extralimital individuals/observations. Additional tables provide the underlying data, compilation parameters, and metadata. The output databases are shareable, compressible, and readable with the R and Python programming languages, and they can be visualized within QGIS projects (QGIS Association 2023) and various graphical user interfaces.

**Table 3.** List of tables included in the output databases generated by the transparent and repeatable range mapping framework. Tables include extensive documentation, and data to provide transparency regarding the basis for maps and decisions that determine resulting maps.

| Table name | Description |
| --- | --- |
| presence | Presence data, including summed occurrence record weights, expert opinion and weight, extralimital status, and presence codes per spatial unit. |
| year_round, summer, or winter | Range data, including summed occurrence record weights, expert opinion and weight, extralimital status, and range codes per spatial unit. |
| simplified_results | Simplified range and presence data per period |
| compilation_info | Metadata and parameters for the range compilation. |
| occurrence_records | The collection of occurrence records that were used in the compilation. |
| opinion_records | The expert opinions that were used in the compilation. |
| range_2001v1 | The GAP version 1 range data for the species (for 2001). |
| last_record | Information about the last occurrence record that was reported within each spatial unit. |
| change_summary | Various metrics of change in presence across time periods. |
| references | Citations for sources of expert opinion. |

**Table 4.** The information hierarchy of the range map compilation process. Documented refers to documentation of presence or range with observational data. Opinions are collected from experts or GAP staff. The Gap Analysis Project published version 1 range maps for 2001 that can be used as an initial map to revise with the compilation process.

| Source 1 | Source 2 | Dominant source |
| --- | --- | --- |
| Documented | Opinion | Documented |
| Opinion | Version 1 | Opinion |
| Opinion | Previous period | Opinion if rank * (confidence/10) > 2 |
| Documented | Version 1 | Documented |
| Documented | Previous period | Documented |
| Previous period | Version 1 | Previous period |

#### Change Summaries

Following the creation and testing of the output database, a summary of changes in species presence across 5-year periods is generated and saved as a JPG. This image file includes a series of panels that visualize:

- Changes in the proportion of the CONUS coded as present and absent for the species.
- Percentage changes in the areas of regions coded as present and absent.
- Proportions of CONUS and total area coded as present where the species was documented.
- Number of GAP HUs where the species was documented compared to the number of HUs with previous documentation.
- Number and mean weight of occurrence records available for the species.
- Ratio of observations to GAP HUs with documented presence.

These trends collectively provide an overview of the stability of species presence maps over time as new data are incorporated. However, increasing trends in data quantity and availability are confounded with actual changes in species’ ranges.

Consequently, the observed trends may only reflect changes in our knowledge of presence rather than true changes in species distributions. Despite this limitation, change summaries visualize how our understanding of species presence evolved over time as new data are incorporated. In some cases, these visualizations may reveal important insights into potential range shifts or other distributional changes.

### Processes

The range compilation process is composed of nine stages. Most are scripted in Python, but some are manual (Figure 1; Table S2).

**Figure 1.**
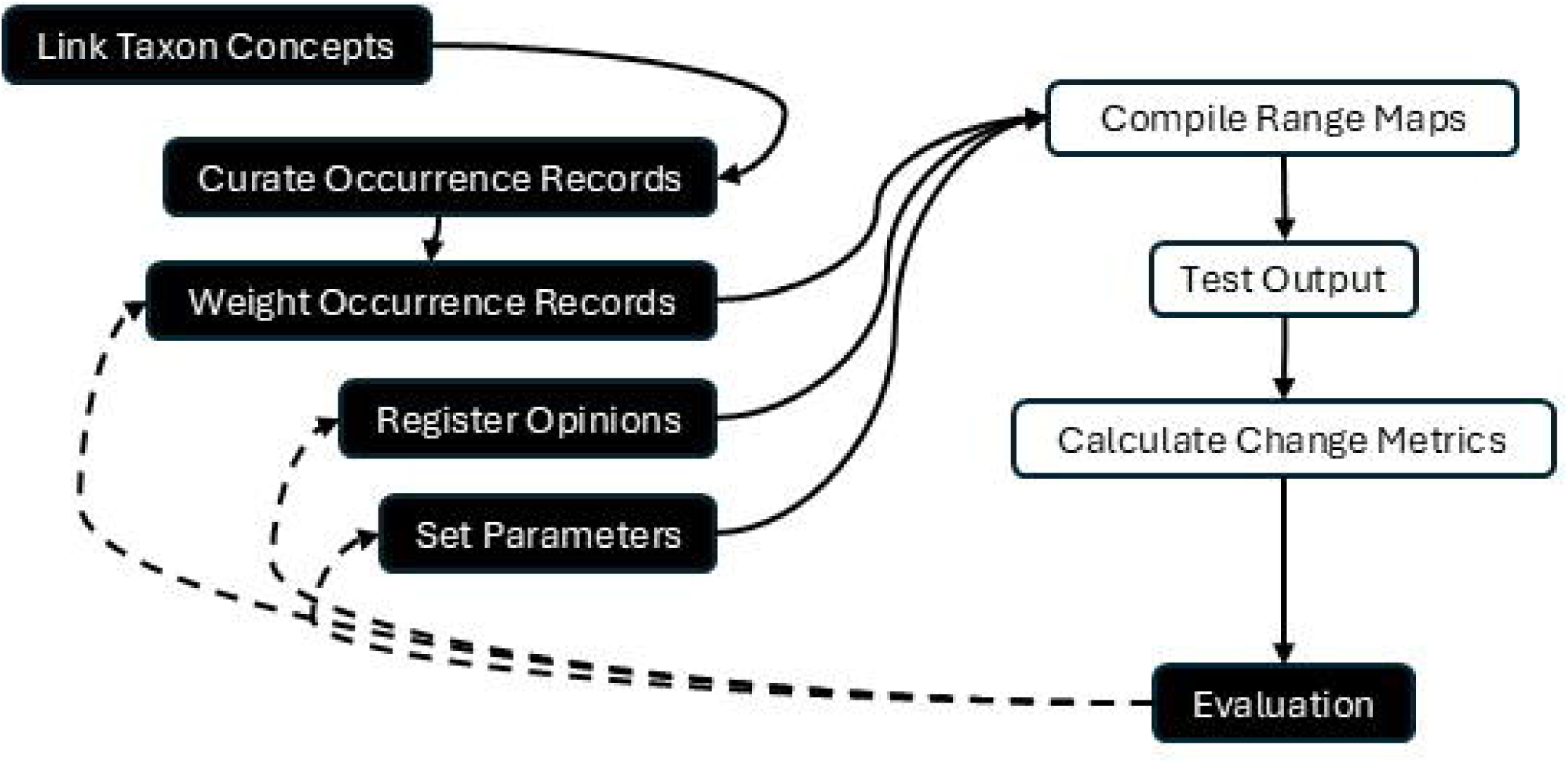
Stages of a species range and presence map compilation framework that integrates different sources of information while providing transparency and repeatability. Arrows indicate the order of sub-processes and dashed lines indicate pathways for refining results. Automated steps are white and manual steps are black. The results of the manual steps are documented extensively in the results.

#### Link Taxon Concepts

The process begins with a critical evaluation of the species names and definitions (taxon concepts) employed by data providers, comparing them to the concepts utilized by GAP. The core objective is to identify all distinct taxon concepts that may exhibit taxonomic overlap with the GAP-accepted concept for the target species, along with their interrelationships over time. This task is important because species occurrence records and expert opinion were assigned to accepted taxon concepts at the time of entry into datasets, which can lead to the incorporation of inappropriate observations or opinions (e.g., for the wrong taxon) that introduce errors into range maps.

Multiple taxon concepts may overlap with a single preferred concept when authorities, data providers, or experts have disagreed about species definitions or nomenclature. Furthermore, past taxonomic revisions may have involved concepts to consider, particularly when incorporating data from previous decades because some data providers may not have updated their records to reflect currently accepted concepts.

While some data infrastructure and software tools exist that can help evaluate relationships between taxa, current resources primarily focus on linking biological nomenclature rather than the underlying taxon concepts themselves. Consequently, concept matching currently necessitates a manual approach. This involves careful consideration of the geographic distribution of individuals implied or identified within each species’ definition.

Manual evaluation of taxon concepts typically involves reviewing online literature and consulting resources, such as NatureServe Explorer (NatureServe 2023) and Avibase (Avibase 2024). The results of these reviews can be stored in a dedicated table where each unique taxon concept is assigned a record with columns delineating the relationships between concepts. The table enables users to navigate taxon overlaps and transitions. Additionally, the table includes scientific and common names, along with identifiers from relevant taxonomic databases (GBIF, Integrated Taxonomic Information System, NatureServe, the Mammal Diversity Database, the American Ornithological Society, and eBird). An optional column is also included for well-known text of a coarse polygon to support delineation of important bounds of the taxon’s geographic distribution for cases where it is beneficial to impose coarse geographic limits on which observations to include.

#### Curate Occurrence Records

While species occurrence data are readily available from GBIF, they often require significant processing to filter out low-quality records. Curation of GBIF datasets can be complex due to the numerous attributes within each record, which provide background information on data sources, collection details, and associated uncertainties. To address this, we developed a separate framework and set of processes for curating species occurrence record datasets, termed the Wildlife Wrangler (Tarr et al., 2022). The Wildlife Wrangler processes GBIF data and produces SQLite databases containing curated occurrence records. The GAP range compiler was designed to utilize occurrence records derived from the Wildlife Wrangler. When multiple taxon concepts have been recognized for a species after 2001, it is necessary to independently process each concept.

Occurrence records are best represented as polygons because they represent events with spatial extents (Chapman and Wieczorek 2020), and the size and shape of record extents are determined by whether the record was recorded as a point or assigned to a sampling unit (e.g. survey block). If recorded as a point, the extent is determined by the path traveled by the observer, the distance at which the species can be detected under the survey methods, GPS accuracy, and the spatial precision of the coordinates (“nominal coordinate precision”). The “CoordinateUncertaintyInMeters” field in GBIF reports these locational uncertainties, but extents are not reported for many records. If desired, the Wildlife Wrangler approximates uncertainty according to methods described by Chapman and Wieczorek (2020). Records can then be represented as polygons during a process of assigning records to spatial units described below.

#### Weight Occurrence Records

Following curation, individual occurrence records are assigned a weight reflecting their perceived quality. This quality assessment is based on attributes extracted or generated by the Wildlife Wrangler. A weight score ranging from zero to ten is assigned to each record. Initially, all records are assigned a maximum weight of ten. Subsequently, the weight is adjusted downward for records exhibiting problematic attributes. For instance, records based on fossil specimens are assigned a weight of zero, because they are considered inappropriate for the temporal extent of GAP data. Weight adjustments are implemented using a Python script, allowing for iterative refinement, re-execution, and archiving of the weight assignment process.

#### Register Opinions

I developed Python scripts that facilitate registration and review of expert opinions using QGIS. To register opinions, the user selects spatial units from a Shapefile of GAP HUs within a QGIS map document and specifies a GAP species code, season (or “presence”), start year and end year, status (present or absent), justification, reference codes that link to entries in the GAP Wildlife Habitat Relationship Database, user initials, and confidence level (1-10). An expert rank is also needed, so the user must self-rank their expertise level (1-10) for the species by spatial unit and year. The script then adds a record to the Opinions Database for each selected spatial unit and year within the specified years.

#### Set Parameters

The user manually enters compilation information (see Table 2) into the range parameters table of the range parameters database.

#### Compile Range Maps

Once the taxon concepts have been resolved and the occurrence and opinion data curated and weighted, the main range compilation processes are run (see Text S2 for details; Figure 2). Range compilation is programmed in a single Python script that utilizes parallel processing to compile presence and seasonal range maps. The compilation script is written as a collection of custom Python functions that each perform a discrete task. Some functions are run in sequence whereas others are run in parallel. Although it is written in Python, the script utilizes SQL programming language statements and SQLite databases “in-memory” for data management and processing. All data are projected to the Alber Equal Area projections for spatial processes (EPSG: 5070).

**Figure 2.**
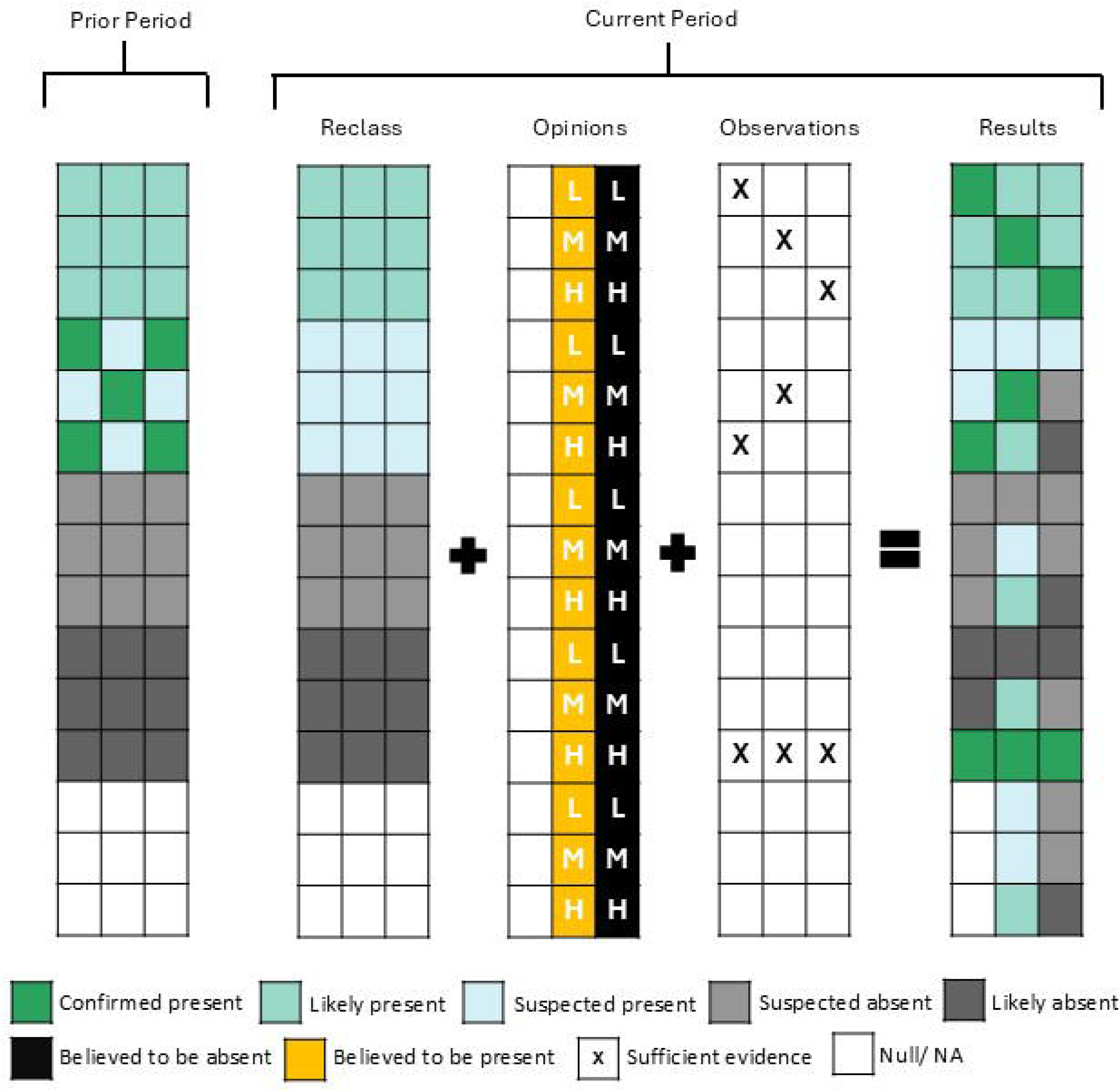
The application of rules for assigning presence codes in a range mapping framework that provides transparency and repeatability. Maps are compiled for discrete time periods of 5 years based upon expert opinion data and species observations for the target (“current”) period and the map from the prior period. Grid cells represent spatial units, and letters in the opinions grid denote opinion weight: “L” for low, “M” for moderate, and “H” for high. Confirmation from the prior period is first reclassified to “likely present”. Then, codes are adjusted where opinion weights are sufficient. Finally, “presence confirmed” is applied where the weight of evidence from observations is sufficient. For the first step (2001-2005), GAP version 1 maps are reclassified and used as the prior period map. An analogous process is applied during range map compilation.

The assignment of occurrence records to spatial units is central to the compilation process. Whether presence and range were documented in each spatial unit is assessed by evaluating the number, weights, and locations of occurrence records in relation to each spatial unit during each period. Presence or range are considered documented within a period when the sum of record weights is greater than 9. That evaluation is complicated by the irregular shapes, sizes, and positions of records and spatial units. Furthermore, a risk of misallocating records to spatial units exists when they are not completely contained by a single unit. I developed a process for managing misallocation risk with an error tolerance parameter that controls which spatial units records are assigned to, which depends upon spatial overlap between records and units (Text S1; Figure S1).

#### Test Output

Following map compilation, a Python script conducts several tests of the results. If a test is failed, a warning is printed to indicate that something went wrong in the automated processes. Tests are described in Text S2.

#### Calculate Change Metrics

The framework supports evaluation of change over time because presence codes are compiled for multiple time periods. A Python script calculates key change metrics, storing them in a “change summary” table. This table serves as the foundation for generating graphs that visualize temporal trends in presence.

#### Evaluate Results

The final step is for the producer to review the results. If adjustments are needed, they can refine results by adjusting occurrence record weights, adding more expert opinions (e.g., tertiary opinions), or altering parameters and rerunning the compiler scripts. Results are recompiled from scratch each time the script is run, which makes results reproducible and supports versioning of results.

### Case Study

The Fisher is a medium-sized carnivorous mammal in the weasel family that occupies forested landscapes in portions of North America (Powell 1981). The American Society of Mammologists moved the species to the genus *Pekania* from the genus *Martes* following phylogenetic analyses by Koepfli et al. (2012; American Society of Mammologists 2024).

The Fisher’s range contracted during the 19^th^ and 20^th^ centuries due to hunting and habitat loss. Fishers were once present extensively in the Sierra Nevada Mountains, Cascades Range, Klamath Mountains, Olympic Peninsula, and Northern Rocky Mountains, but populations in those areas are now patchy and small (Lewis et al. 2022). The southern Sierra Nevada populations are small and genetically isolated (Tucker 2013; Sweitzer 2021), thus they were recently listed as endangered under US Endangered Species Act (ESA 1973, as amended; Lewis et al. 2022). Washington added the species to its state endangered species list (Lewis et al. 2022). In the eastern United States, Fishers were once present in the Northeast, Upper Midwest, and portions of the Appalachian Mountains as far south as North Carolina and Tennessee, but they were extirpated from many of those areas (Moncrief and Fies 2015).

In recent decades, the Fisher range has expanded because populations have recolonized some areas and conservation programs have translocated individuals into others (Lewis et al. 2012; Lewis et al. 2022). Following extirpation from Washington State in the 1930’s, Fishers were successfully reintroduced to the Olympic Peninsula and Cascade Range beginning in 2008 (Happe et al. 2020; National Park Service 2021; Washington State Washington State Department of Fish and Wildlife 2021; Lewis et al. 2022). Fishers were also believed to have been extirpated from the northern Rocky Mountains, so Idaho and Montana game agencies performed reintroductions in the 1950’s, 60’s, and 90’s (Waller 2018). However, undetected populations had persisted in west-central Montana, despite beliefs that they had been completely extirpated (Lucid et al. 2019; Coltrane and Inman 2021). Fishers have recently recolonized portions of their historic range in the East (Krohner et al. 2021); they were observed in western Maryland and Virginia in recent years, which were the southernmost eastern populations (Moncrief and Fies 2015; Yalcinkaya 2021).

I compiled maps with an error tolerance of 20% and extralimital cutoff at 100 km (Data S1). Processing time was approximately two minutes. The resulting maps consisted of patches of range in the Northeast, Appalachian Mountains, east-central Tennessee, the Upper Midwest, the northern Rocky Mountains, the northern and central Cascades, the Olympic Peninsula, Pacific Coast Ranges, and southern and northern Sierra Nevada Mountains (Figures 3 and S6). In addition, Fishers were believed to be present in portions of Idaho, Wyoming, and Montana, but the records were likely of transient, dispersing individuals that are not indicative of the species’ range (Figures 3 and S5).

**Figure 3.**
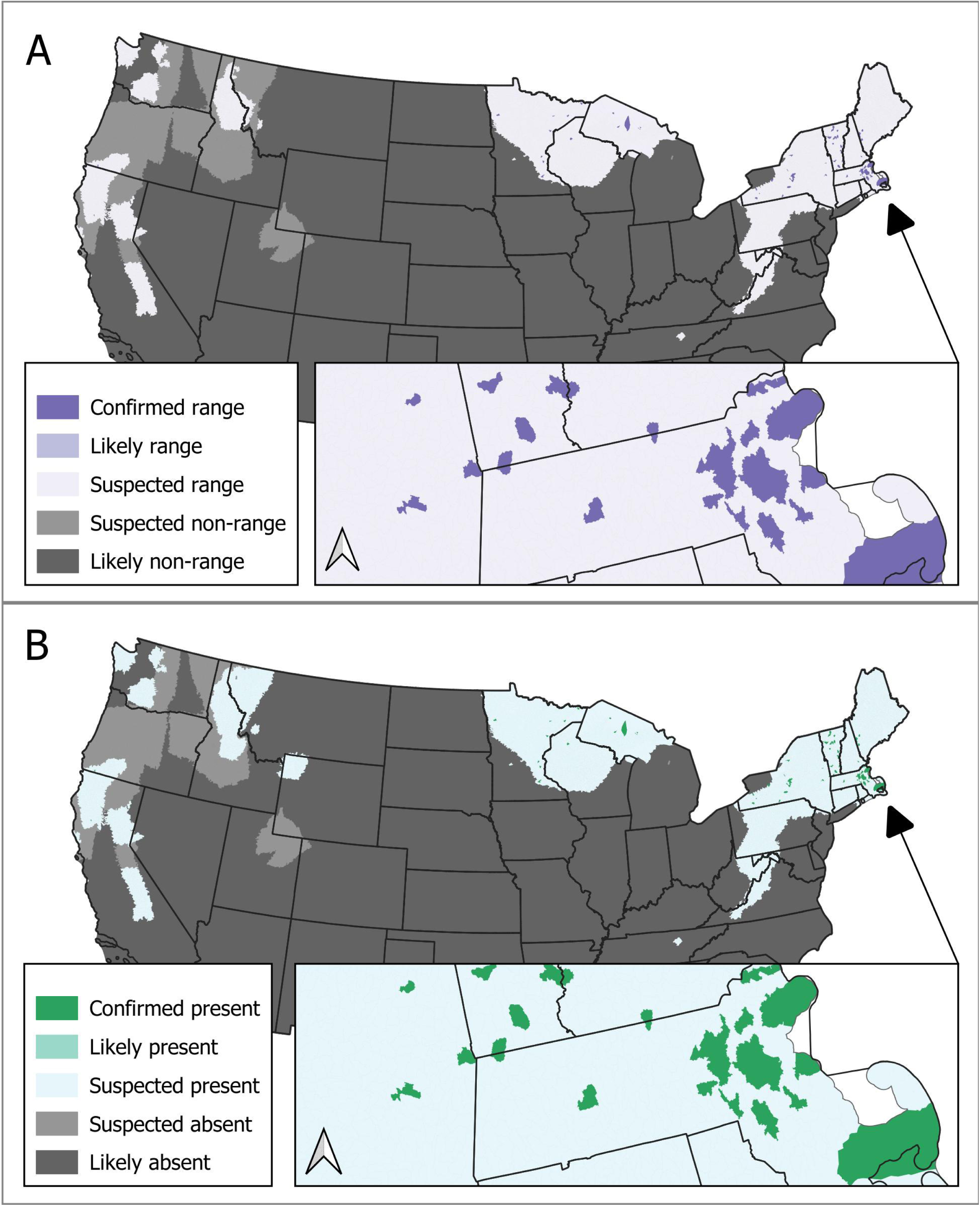
Fisher range (A) and presence (B) within CONUS from 2021 to 2025. The maps were created with a transparent and repeatable framework that integrates information from an existing map, species observations, and expert opinion. Map categories reflect levels of evidence regarding the existence of individuals or range within spatial units. Categories were assigned based upon an information hierarchy, with observations driving confirmation of presence or range. Suspected categories were supported by prior maps and expert opinions. Likely range and presence categories do not occur in these maps and are largely reserved for future efforts to directly incorporate model predictions, although some conditions could generate them in the current version of the framework.

### Fisher Data

#### Initial GAP Range Map

GAP version 1 year-round range and presence maps were identical and depicted the species in the Appalachian Mountains north of North Carolina and south of Maryland; northern Maine and other New England states within ∼100 km of the Canada border; areas immediately surrounding Sault Saint Marie, MI; an area encompassing northern portions of Michigan and Wisconsin, as well as the western portion of Michigan’s Upper Peninsula; a portion of southwest Wyoming and northeastern Utah; the Northern Rockies; the entire Cascade Range; the entire Sierra Nevada Range; and coastal and near-coast areas of northern California and southern Oregon. Version 1 range data indicated the species was absent from Tennessee, Pennsylvania, Maryland, New Jersey, Massachusetts, Rhode Island, and Connecticut; central portions of the Upper Peninsula of Michigan; and the Olympic Peninsula.

#### Fisher Occurrence Records

I queried the GBIF API for Fisher occurrence records on January 23, 2025 and requested data be returned as Darwin Core Archives. The taxonomic revision that moved the Fisher from the genus *Martes* to *Pekania* meant that relevant records could have been associated with either of two taxon concepts in GBIF for our period of interest (2001-2025). Therefore, I queried records with each GBIF taxon identifier (Table S3). For *M. pennanti*, I requested records from 2000 – 2014 because taxonomic revision was around 2014. For *P. pennanti*, I queried records from after 2000.

I accepted records from any month, basis of record, collection, dataset, and sampling protocol but excluded non-United States records because GAP’s current study extent is CONUS. I accepted records without a coordinate uncertainty value but excluded records with a reported coordinate uncertainty greater than 100 km. When no coordinate uncertainty was reported, I approximated uncertainty with the point-radius approach described in Text S2 and Chapman and Wieczorek (2020). I filtered out records with duplicate date-location combinations, throwing out all but one record in each case. I omitted records with flags of “RECORDED_DATE_INVALID”, “TAXON_MATCH_HIGHERRANK”, and “TYPE_STATUS_INVALID” because invalid dates could cause erroneous removal of duplicates and errors in assigning records to periods; taxon matching needs to be correct so that other species’ records are not included; and invalid type statuses could allow the inclusion of fossils. I also omitted twelve records with event date values that were insufficiently precise for our application.

I weighted occurrence records on a scale of 1 to 10 according to uncertainty about the time, location, or identification of organisms. Records for which uncertainty was high were weighted low. Pacific marten, American mink, and river otters can be misidentified as Fishers based on direct observations of individuals or their tracks (Aubry et al. 2017), even by professional biologists (Waller 2018). Therefore, I initially weighted all records at a value of 5 and then adjusted values up or down according to record attributes. With this approach, two or more records were required to document presence within a spatial unit unless a record could be assigned the maximum weight of ten. The following steps were performed to adjust the weights:

- Weights were set to ten if the record’s basis was a preserved specimen, because those records were unlikely to be misidentified.
- Weights were reduced by three if records were flagged as having invalid or unknown geodetic datums, or if longitude values were presumed to have been negated, because such flags indicated uncertainty and imprecision in the locations.
- Weights were set to zero for records flagged with invalid dates or geographic coordinates that may have been swapped because those records could introduce large errors in range limits.
- Weights were also set to zero if remarks about a record’s location indicated the provided coordinates were in the center of a state, county, or other broad region because the spatial precision of those records was insufficient for our application.
- We set the weight of record 2564830634 (see gbif.org/occurrence/2564830634) to zero because the occurrence remarks contradicted the coordinate uncertainty value.
- We decreased the weight by five for records 3079750531, 3903399252, 3090747664, 3966597768, and 3784795964 because the included remarks indicated uncertainty about the identification of the individual.

The filters allowed the retention of 2,287 of 2,444 available *P. pennanti* records (GBIF 2025a; File S1) and 148 of 175 available *M. pennanti* records (GBIF 2025b; File S2). For *M. pennanti*, 20 duplicate records were dropped and 50 had coordinate uncertainty values. For *P. pennanti*, 138 duplicates were dropped and 1,460 records had uncertainty values. Upon review of the results (GBIF 2025a; GBIF 2025b) we determined that all *M. pennanti* records were included in the *P. pennanti* records as well. Therefore, we discarded the *M. pennanti* query results and relied solely on the 2,287 *P. pennanti* records.

1. *P. pennanti* records were available from 16 datasets (Text S3). The iNaturalist Research-grade Observations dataset included the most records (n = 1,584, iNaturalist contributors 2025), followed by the KUBI Mammalogy Collection (n = 604, Bentley and Krejsa 2024) and others (see Text S3 or GBIF 2025a for a complete list). All records were based on human observation or preserved specimens. No sampling protocol was reported for 2,429 records but the remaining records were from trapping, shooting, or road killed. Ten issue flags were present in the dataset, which we used to inform record weighting:
2. “COORDINATE_ROUNDED”
3. “COLLECTION_MATCH_FUZZY”
4. “INSTITUTION_COLLECTION_MISMATCH”
5. “INSTITUTION_MATCH_FUZZY”
6. “GEODETIC_DATUM_ASSUMED_WGS84”
7. “COORDINATE_REPROJECTED”
8. “OCCURRENCE_STATUS_INFERRED_FROM_INDIVIDUAL_COUNT”
9. “TAXON_MATCH_TAXON_ID_IGNORED”
10. “GEODETIC_DATUM_INVALID”
11. “CONTINENT_DERIVED_FROM_COORDINATES”

Occurrence records were available from all large patches of range except for the Northern Rockies, Tennessee, and the Northern Cascades (Figure 4). Many of the records from the Upper Peninsula of Michigan were decades old (File S1, Figure S4), but recent records were available for many regions. The distribution of records by year was bimodal with few records available from 2005 to 2015 (File S1). Most records from before 2005 were from the Upper Peninsula (Table S8). Records were also unevenly distributed across months (File S1), with more winter than summer records and an inordinate number of records from December, which could be attributed to records from the Upper Peninsula that were provided to the KUBI Mammalogy Collection by the Michigan Department of Natural Resources.

**Figure 4.**
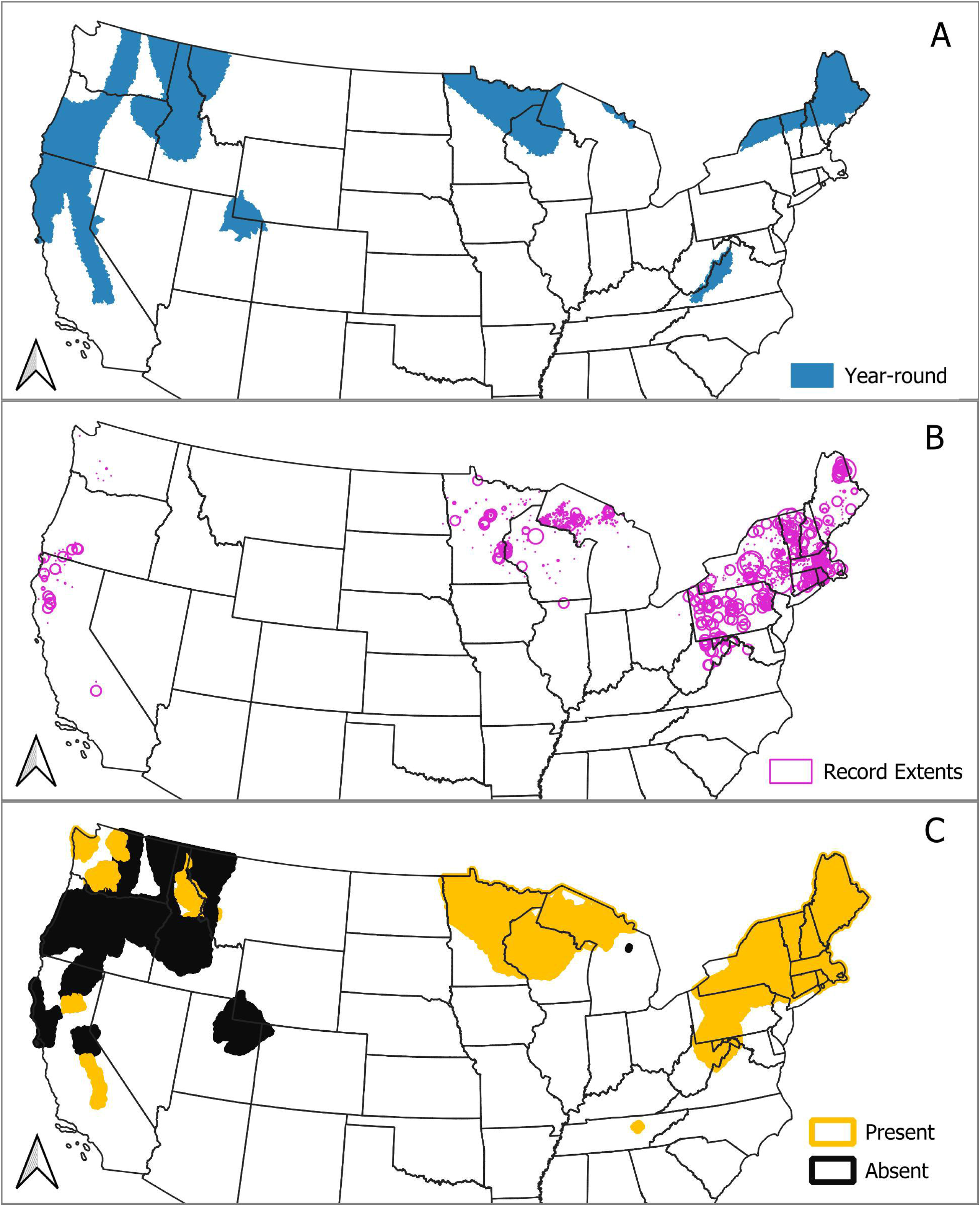
Data used for a Fisher range map within CONUS from 2021 to 2025. The compilation process integrated a prior map (A), species occurrence records (B), and expert opinion regarding the presence of range (C) according to standardized rules. The GAP version 1 Fisher range map for 2001 provided an initial map (A). Fisher occurrence record extents (2001-2025), which are geographic coordinates buffered with locational uncertainty supported the addition of some spatial units (B). Expert opinion regarding Fisher range (2001-2025) also supported the addition and removal of spatial units (C).

Nominal coordinate precisions were mostly below 12 m, but some were over 1,000 m (File S1). The median point-radius buffer distance was 188 m, but locational uncertainty was large for many records: we computed point-buffer radii larger than 2 km for 441 records; File S1). The records with the largest locational uncertainty came from the iNaturalist Research-grade Observations dataset (iNaturalist contributors 2025).

We adjusted the buffer radius of some records for which coordinate uncertainty was reported to be zero. We found more information about the records on the Kansas University Biodiversity Institute Mammalogy Collection web portal and determined that the coordinate uncertainties were actually 1,191 m or 1,030 m (Bentley and Krejsa 2024).

Of the 2,287 Fisher occurrence records that we acquired, 14 were not used because their weights were adjusted to zero. All other records were given weights of 10 (n = 597), 5 (n = 1,674), or 4 (n = 2). Mean record weight declined from 2001 to 2025 (Figure S5).

#### Expert Opinions

I incorporated secondary expert opinions on Fisher range and presence from 16 sources (Text S4; Figure 4). In combination with tertiary opinions, I found justification for the addition and removal of large areas from range maps in the 2016-2020 (Figure S6) and the 2001-2005 steps (Figure S6). Expert opinion reflected belief that the species was present in New England; the Upper Midwest; mountainous portions of Pennsylvania, Maryland, and West Virginia; a portion of Tennessee; portions of northwestern Washington including the Olympic Peninsula; portions of the Sierra Nevada Range and Northern Rocky Mountains. Absence was indicated for Utah and southwestern Wyoming; the Sierra Nevada Mountains between Yosemite National Park and Carson City, NV; the California coast between Eureka and San Francisco; portions of the Cascade Range; portions of the Northern Rocky Mountains; and nearly all of Oregon except for portions of the Klamath Mountains. Regarding Fisher presence, expert opinion aligned with beliefs about range, except that non-range presence was indicated for northwestern Wyoming and western Montana near the Idaho border (Figure S7).

## Discussion

This framework facilitates the revision of species range maps by integrating species observations and expert opinions. It adheres to explicit definitions of range and standardized compilation methods to ensure clear, repeatable, and reproducible range maps. The framework overcomes several challenges inherent to mapping ranges and provides a practical method for maintaining, updating, and revising maps as information and species distributions change over time.

- The framework can track changes in species distributions, including shifts, retractions, and perforations. This capability stems from the explicit definition of spatiotemporal units and the inclusion of absence in the map legends. These features also contribute to improved precision in delineating patches of range.
- The framework provides transparency because underlying data are preserved and included alongside results, many processes are automated and controlled by explicit parameters, and non-automated content can be thoroughly documented. Transparency means that users and reviewers can clearly interpret the data and revisions can build upon previous versions instead of “starting from scratch”.
- The structure of the framework, especially the scripted portions, ensure repeatability and iterative refinement of maps, which is central to the review and revision of maps. Furthermore, the framework is compatible with map versioning, which can improve data provenance and help organize the revision process.
- The framework reduces speculation about species’ ranges by map creators and incorporates documentation that reveals experts’ confidence in opinions.
- The framework addresses risks of allocating observations to the wrong spatial unit to avoid errors of commission, which are especially undesirable and problematic for GAP’s habitat mapping framework. The method is powerful when combined with occurrence record weights because it accounts for sources of uncertainty beyond spatial precision whereby low-weighted records with low spatial precision have minimal influence on the results.
- By relying upon explicit taxon concepts as opposed to taxon names only, the framework avoids confusion regarding the specific taxon represented in the maps. This approach also reduces the risk of commission errors from inappropriate occurrence records and omission of valuable occurrence records.
- The framework leverages open-source software with parallel processing capabilities. This significantly reduces the human workload by automating time-consuming tasks. For example, runtimes for certain processes can be reduced from hours or days to minutes or seconds. This frees up staff time to focus on tasks that require human expertise and judgment.
- The framework acknowledges data providers by retrieving and including citations for observations and references. Similarly, publications and experts that provide information are clearly cited.

### Fisher Case Study

I produced Fisher range and presence maps for the period 2021 - 2025 by revising existing data based upon Fisher occurrence records and expert opinions. The new maps exclude large unoccupied areas that were included in GAP version 1 maps. Other areas were added to reflect range expansions that were initiated by translocations of individuals by national, tribal, and state governments along with non-governmental conservations groups. Our maps appear to have greater precision than current IUCN (IUCN 2016) and Map of Life (2021) Fisher range maps and identify previously unincorporated areas of range.

The maps could support the addition and removal of Fisher from several state species lists. The information we reviewed indicated that Fisher range does not extend into Utah or Wyoming. Additionally, Fisher range is less extensive in the Northwest than indicated by GAP version 1. However, the eastern range appears to have expanded south towards the southern Appalachian Mountains. If this expansion continues, North Carolina, Georgia, and South Carolina may need to consider adding the Fisher to state species lists.

We incorporated over 1,000 records from museums and community scientists into the compilation process and we were able to cite data providers and provide transparency for our results. Accessing occurrence data through GBIF provided efficient ways to manage and assess records in bulk, but also provided the ability to thoroughly examine individual records on the GBIF website when necessary. Utilizing the Wildlife Wrangler allowed us to curate and archive an observation dataset that included all relevant attributes to inform the maps. The maps benefited greatly from data provided by Michigan Department of Natural Resources. They provided many high-quality records that revealed the species’ presence in the Upper Peninsula of Michigan.

The advent of iNaturalist greatly increased the number of records that were available, but the spatial precision of iNaturalist records proved to be a challenge: many records had locational uncertainty that was too great to support attribution to one of our spatial units. Additionally, I weighted over 1,000 records too low to confirm presence or range without presence of another record because I could not verify their legitimacy. However, it is likely possible to utilize more of those records with more precise and accurate methods for identifying high-quality records.

Numerous peer-reviewed publications are available that describe at least some portion of the Fisher’s range (Text S3). Several of the publications detail reintroductions or predictive models from the last decade that reflect recent funding for Fisher conservation (Coltrane and Inman 2021; Green et al. 2022; Lucid et al. 2021). Others described or investigated occurrences in notable locations (Waller 2018; Moncrief and Fies 2015). Those publications sometimes included information about the Fisher’s range during prior decades, in addition to information from their study periods. In such cases, it was possible to incorporate both types of information into the maps with the expert opinion framework.

### Potential Improvements and Extensions

The framework is presented as a practical baseline that can be improved over time by addressing key challenges:

#### Efficiency in Resolving Taxonomy

The process of identifying and matching taxon concepts can be very time consuming. Future reviews can build off previous work, but taxonomy needs to be reviewed periodically to identify and account for taxonomic revisions by authorities. Automated processes that rely upon concept-based linkages could speed up the process. The Biodiversity Information Standards Taxon Concept Schema Maintenance Group (TDWG 2024) has begun developing a data standard that may prove useful.

#### Evaluating Occurrence Records

Ideally, all occurrence records would undergo individual examination to identify errors and apply appropriate weights. However, manual review is extremely time-consuming. Rego et al. (2024) individually reviewed 620,000 records for 4,148 Amazonian bird taxa and that task required labor equivalent to approximately two and a half to three years of work by one person.

Labor constraints necessitate approaches that remove and weight records in broad strokes, which may exclude large proportions of records. For example, I omitted dozens of records for the Fisher and weighted over a thousand low enough that they had minimal influence on results. Machine learning algorithms could be explored as a means for more efficient and accurate weighting according to project objectives.

Additionally, some species occurrence records that have been assigned an error flag by GBIF could be corrected (i.e., “cleaned”) instead of being addressed with filters or low weights. I did not include a cleaning step in this iteration of the framework, but cleaning could be performed between the curation and weighting of occurrence datasets (Figure 1). Correcting low-quality data and boosting their weights would increase the number of records used. It may be possible to develop R scripts that utilize the CoordinateCleaner (Zizka et al. 2019) package for this task.

#### Identifying Extralimital Records

The identification of extralimital records is inherently difficult and a valuable topic for further development. Two challenges are worth considering, and although they may be unavoidable, they can be addressed with documentation provided by this framework. One, occurrence records need to be compared to a range limit to determine whether they are extralimital, but extralimital records need to be ignored when delineating those range limits. Hence, knowledge of range limits is paradoxically necessary for the data-driven delineation of range limits. Two, data-driven range delineation methods must be sensitive to signals of colonization yet resistant to the influence of extralimital records. In other words, records that represent true range or colonization events are at risk of being flagged as extralimital.

Potential criteria for identifying extralimital records are imperfect. For example, a threshold distance from other records or hypothesized range (e.g., the “extralimital cutoff” implemented in the framework) can be applied beyond which records are considered extralimital. While this method can effectively flag long-distance dispersers and migrants that have strayed far from their expected range, it may exclude records from newly colonized or previously unknown disjunct populations. Moreover, determining an appropriate distance threshold remains an ambiguous task.

Alternatively, a threshold based on the number of records within a given area could be applied, with areas containing fewer than a specified number of records getting classified as extralimital. In this approach, sparse or inconsistent spatiotemporal coverage of occurrence datasets may lead to the exclusion of valuable intralimital records.

Furthermore, the rarity of certain species often attracts increased observation effort, particularly for avian taxa (Lees and Gilroy 2021). This can result in a large number of records for a single or few individuals at a specific location, potentially leading to their misclassification as intralimital even if they occur within the species’ range. To address this issue, a frequency of observation threshold could be implemented. However, that approach is also susceptible to biases in sampling effort and gaps in occurrence data, potentially leading to the exclusion of genuine intralimital records.

#### Incorporation of Model Predictions

The framework was designed to incorporate output from predictive models directly within the information hierarchy and legends. This would be achieved by utilizing the “likely” categories to allow model predictions to potentially override expert opinions or prior map information. However, I have not yet developed a mechanism for the direct integration of predictive model results; they are currently incorporated as secondary expert opinions, as exemplified by the Fisher case study where we referenced Krohner et al. (2022). This approach limits the impact of predictive models, as they can only trigger “suspected” presence codes when entered as secondary opinion. To fully accommodate the value of predictive models, it would be desirable to develop a pathway for directly incorporating their outputs to populate spatial units with the “likely” codes.

#### Expert Opinions

Expert opinion is central to the creation of range maps because the spatial and temporal coverage of observational data and reliable model predictions is incomplete for many species. Our framework supports the collection, organization, and synthesis of primary, secondary, and tertiary opinions, but we have yet to pursue experts to provide primary opinions. Primary opinions could be collected by asking experts to submit opinions (present/absent or range/non-range) with associated confidence and justification for specific spatial units during specified years.

### Extensions

The framework could be transferred to other regions or projects with a few changes. One, a grid of spatial units to build upon must be designated and created as an SQLite database. Two, methods for resolving taxonomic matches must be identified, although the method presented here would likely suffice. Three, an expert opinion database is needed that links opinions to the project’s spatial units. Four, a parameters database needs to be created, although that database is essentially a single table as described above in SQLite format.

This framework could facilitate public participation in improvements to range maps because it builds upon publicly accessible infrastructure developed and maintained by GBIF. The Global Biodiversity Information Facility serves data from many providers, some of which gather data from community scientists and amateur naturalists, such as iNaturalist and eBird. Thus, contributions to those projects quickly become available for applications of this framework.

### Conclusion

This framework provides a transparent methodology for revising species range maps with open-source tools and the integration of diverse data. While the Fisher case study demonstrates the framework’s efficacy in producing more accurate and refined maps, potential improvements and extensions, such as automated evaluation of occurrence data and direct incorporation of predictive models, offer avenues for further refinement.

Ultimately, this framework provides a practical and adaptable foundation for maintaining and updating species range maps, ensuring they remain relevant and accurate in the face of new information and changing species distributions.

## Supplemental Materials

**Text S1.** Additional background on approaches to mapping species ranges.

**Text S2.** Detailed text describing the automated range compilation processes, as well as the tests applied to results. A Python script integrates Gap Analysis Project version 1 range data for 2001, species occurrence records from the Global Biodiversity Information Facility, and expert opinion into range and presence maps according to standardized rules and flexible parameters. The script uses parallel processing and SQLite for efficiency.

**Text S3.** References for the 16 species occurrence datasets from the Global Biodiversity Information Facility that provided records for a Fisher *Pekania pennanti* range map. I applied a transparent and repeatable range mapping framework that integrated occurrence records and expert opinions to update maps for CONUS between 2021 and 2025.

**Text S4.** Annotated bibliography of sources for secondary expert opinion regarding the distribution of Fishers *Pekania pennanti* between 2001 and 2025 in CONUS. In the transparent and repeatable range mapping framework, expert opinion is defined loosely and three types are recognized. Primary opinions comes from biologists and are based upon their direct knowledge and experience. Secondary opinions are gleaned from scientific publications, and tertiary opinions come from map producers’ comprehensive review of available information.

**File S1.** Wildlife Wrangler output report for *Pekania pennanti*. The Wildlife Wrangler is a software tool that facilitates transparent and repeatable queries of GBIF for species occurrence records and includes filters to remove undesirable records. Users interact with the Wildlife Wrangler with a Jupyter Notebook document that includes descriptions of the occurrence records retrieved and is exported as an html file for documentation. The contents of html documents can be viewed with a web browser.

**File S2.** Wildlife Wrangler output report for *Martes pennanti*. The Wildlife Wrangler is a software tool that facilitates transparent and repeatable queries of GBIF for species occurrence records and includes filters to remove undesirable records. Users interact with the Wildlife Wrangler with a Jupyter Notebook document that includes descriptions of the occurrence records retrieved and is exported as an html file for documentation. The contents of html documents can be viewed with a web browser.

**Data S1.** SQLite database of the compiled range dataset, underlying data, and documentation. The database includes spatial metadata and can be read with programming languages, such as R and Python; viewed in QGIS and other geographic information systems; and accessed with various graphical user interfaces.

**Table S1.** Column descriptions for presence, summer, winter, and year-round tables in an expert opinion database that provides data for the transparent and repeatable framework for range map compilation.

**Table S2.** Associations between compilation steps and databases in the transparent and repeatable range framework.

**Table S3.** Taxon concepts included in queries of the Global Biodiversity Information Facility for Fisher *Pekania pennanti* occurrence records to support the compilation of an updated range map. “GAP ID” is the National Gap Analysis Project species code, “GBIF ID” is the Global Biodiversity Information Facility’s identification code, “ITIS TSN” is the Integrated Taxonomic Information System’s Taxonomic Serial Number, “MDD ID” is the American Society for Mammologist’s Mammal Diversity Database taxon identification code, and “NatureServe ID” is the suffix of the NatureServe Unique Identifier.

**Figure S1.** Examples of allocation error risks associated with occurrence records overlapping multiple spatial units. To incorporate occurrence data into gridded range maps, records must be assigned to spatial units. Risks of allocation errors arise when the geographic extents of records are not completely contained within spatial units. Squares represent gridded spatial units and circles represent species occurrence record polygons. The size and position of occurrence records is identical in each panel and allocation error risk is reported for each spatial unit. Each panel shows the consequences of a different level of tolerance for allocation errors. Grey spatial units are those that would be attributed to the occurrence record under the specified error tolerance.

**Figure S2.** Fisher *Pekania pennanti* range maps for each 5-year period from 2001 to 2025 generated with a framework that integrates underlying data in time steps. Changes over time periods reflect changes in the availability of information, as well as actual expansions and contractions in the geographic distribution of the species.

**Figure S3.** Fisher *Pekania pennanti* presence maps for each 5-year period from 2001 to 2025 generated with a framework that integrates underlying data in time steps. Changes over time periods reflect changes in the availability of information, as well as actual expansions and contractions in the geographic distribution of the species.

**Figure S4.** Age in weeks of the most recent Fisher *Pekania pennanti* record attributed to spatial units within the conterminous United States as of January 23, 2025. I downloaded the occurrence records from the Global Biodiversity Information Facility.

**Figure S5.** Metrics of change in Fisher *Pekania pennanti* presence across time periods. The metrics were generated with a framework that integrates underlying data for time steps. Changes over time periods reflect changes in the availability of information, as well as actual expansions and contractions in the geographic distribution of the species.

**Figure S6.** Secondary and tertiary expert opinions about Fisher *Pekania pennanti* range aggregated by period. Opinions were registered by year and spatial unit in a private database. Secondary opinions were gleaned from scientific literature and tertiary opinions were formed during the review of compilation results at intermediate stages.

**Figure S7.** Secondary and tertiary expert opinions about Fisher *Pekania pennanti* presence aggregated by compilation period. Opinions were registered by year and spatial unit in a private database. Secondary opinions were gleaned from scientific literature and tertiary opinions were formed during the review of compilation results at intermediate stages.

**Figure S8.** Illustration of a case where “suspected presence” (or “suspected range”) codes would be adjusted to “likely present” in a range mapping framework in which range maps are compiled in 5-year increments for spatial units. All boxes represent information for a single spatial unit.

## Supporting information

Project Data

Occurrence Record Queries

## Acknowledgements

The U.S. Geological Survey (USGS) Science Synthesis, Analysis and Research Program (M. Wiltermuth, Project Manager) provided funding for this research under Research Work Order 236 with the North Carolina Cooperative Fish and Wildlife Research Unit. I thank Alexa McKerrow, Ken Boykin, Steve Williams, Matthew Rubino, Nathan Hostetter, Mark Wiltermuth, Curtis Belyea, Anne Davidson, and Leah Dunn for feedback on the framework and early versions of the manuscript. I also thank GBIF staff, data providers, and the biodiversity data infrastructure community for enabling the application of species observations to conservation biogeography. Google Gemini was used to improve the clarity and brevity of portions of text, but the author reviewed and edited the results and is responsible for the content.

