## Supplemental Tables for "A Framework for Transparent and Repeatable Species Range Maps"

Table S1.

| **Column** | **Description** |
| --- | --- |
| HUC | The Gap Analysis Project’s unique code for the spatial unit. |
| Species | The unique Gap Analysis Project species code. |
| Status | Whether the species or range is present or absent in the spatial unit. |
| Confidence | Expert self-ranked confidence on a scale of 1 to 10. |
| Year | The year that the opinion applies to. |
| Justification | A detailed justification for the opinion about the status, including citations. |
| Citations | A list of citation codes that link to full citation strings in a private Gap Analysis Project database. |
| Expert | The expert’s name. |
| Expert rank | A rank from 0 to 10 assigned to the expert by Gap Analysis Project staff. |
| Entry time | The date and time when the opinion was added to the database. |

Table S2.

| **Processing Step** | **Database** | **Action** |
| --- | --- | --- |
| Link taxon concepts | Taxa Map Database | Fill out and/or revise records |
| Curate occurrence records | Wildlife Wrangler output database(s) | Write to database |
| Weight occurrence records | Wildlife Wrangler output database(s) | Update values in database |
| Register opinions | Opinions database | Write to database |
| Set parameters | Parameters database | Write to database |
| Compile range maps | Opinions, parameters, occurrence records; range output database | Read from databases and write to output database |
| Test output | Range output database | Read from database |
| Calculate change metrics | Range output database | Read from database, write to database |
| Evaluation | Range output database | Read from database |

Table S3.

| Concept | Scientific Name | GAP ID | GBIF ID | ITIS TSN | MDD ID | NatureServe ID |
| --- | --- | --- | --- | --- | --- | --- |
| A | *Pekania pennanti* | mFISHx | 8631083 | 1086061 | 1005825 | 103714 |
| B | *Martes pennanti* | - | 5218855 | 180560 | - | - |
