## Supplemental Text 1 for "A Framework for Transparent and Repeatable Species Range Maps"

**Methods for Mapping Ranges**

Range maps would ideally be compiled from complete data on the areas occupied by individuals of a taxon within a defined period, preferably with a clear basis for designating individuals as geographic outliers. Range mapping would then only entail delineating a boundary around the areas used by non-outliers (extralimitals). Unsurprisingly, such occurrence data and clarity regarding which records are extralimital are unavailable for most species. Instead, range mapping requires the acquisition, synthesis, and utilization of different types of data and information, each of which is usually imperfect and incomplete (Jetz et al. 2019, Marsh et al. 2022). Thus, range maps are the products of human assessments and interpretations of disparate pieces of information. Data gaps can constrain and define the mapping process. Hopefully, the range compilation process produces an approximation of true range that is accurate enough to support the intended applications. Inaccuracies would take the form of either commission or omission errors (Guisan et al. 2017).

Species occurrence records are often utilized. Public repositories of data, such as the Global Biodiversity Information Facility (GBIF), are now readily accessible via websites and APIs (GBIF 2023). For some species, additional high-quality datasets may exist that have not been shared publicly, although access to such datasets can be limited. Species occurrence records that are deemed reliable can reveal areas where a species was known to occur. Furthermore, records can document or confirm range after extralimital records are identified and removed. Intralimital records can also be used as a basis for delineating range boundaries (Graham and Hijmans 2006) and as training data for predictive models of occurrence (Guisan et al. 2017).

Experts may have experience and knowledge regarding species ranges that differ from what is captured by species occurrence records and associated models. Thus, experts’ opinions about whether areas are within or outside of range limits can fill gaps in other data sources (Marsh et al. 2022). Opinions about absence can be especially valuable for revising range maps because of the difficulty in establishing absence with empirical data (Jetz et al. 2019). However, opinions may also be inaccurate.

Expert opinion can be directly solicited and integrated as polygon or point records, but it can also be gleaned from textual descriptions of species’ geographies in scientific literature, such as journal articles and species descriptions (Jetz et al. 2019). Utilizing those resources requires an added step of converting the knowledge to a spatial data format. Range maps that were created in the past by experts with extensive knowledge of species’ distributions or who previously synthesized information on the species range may also be available. Existing maps are not always available in a georeferenced format, but methods for converting them into spatial data formats are available (Marsh et al. 2022).

It is worth noting that although they are different types of data, occurrence records and opinions are not independent of each other during range compilation. Expert opinion plays a role in the assessment and application of occurrence records and in the design of predictive models. Conversely, readily available occurrence records may also affect the opinions and beliefs of experts before they are solicited. These mutual influences can create challenges during attempts to apply transparent, data-driven approaches to range mapping.

With data in hand, range mapping is fundamentally an exercise in delineation and interpolation whereby the goal is to reveal the geographic limits of the areas where most individuals occurred during a time period. Range maps can be in the form of a single or few polygons in a vector file, attribution to a systematic or natural grid of polygons in a vector file, or predictions applied to a raster grid. The delineation aspect is more obvious when working with un-gridded polygons in a vector format, but mapping with raster or vector grids arguably involves delineation as well, because the emphasis remains on revealing the geographic limits of range.

Several methods can be used for delineating ranges. Experts or non-experts can hand-draw polygons to represent range limits where they believe them to exist, based upon available information and individual knowledge and experience (Graham and Hijmans 2006, Jetz et al. 2019, Marsh et al. 2022). Alternatively, polygons can be drawn around the locations of species occurrence records manually or with computational geometry, such as alpha-hulls (IUCN 2022, Rego et al. 2024). A “point-to-grid” approach can be taken in which occurrence record locations are intersected with a grid to identify spatial units containing records (Graham and Hijmans 2006). Occurrence record locations could instead be buffered to a specified distance to approximate range limits (Aubry et al. 2017, Mims et al. 2018). Range limits have also been approximated with statistical methods that incorporated population growth and dispersal models to predict the spread of populations from known locations over time (Barry et al. 2021). Finally, sampling designs that support occupancy models can be utilized to refine coarse understandings of range limits by removing areas with insufficient probability of occurrence (Krohner et al. 2022). With any of these methods, errors are introduced when gaps and biases exist in the data, but the point-to-grid and point buffering approaches are especially vulnerable to such errors when few occurrence records are available.

The spatial coverages of data and knowledge are rarely complete and often biased, thus it is often necessary to interpolate between areas of known range to fill gaps (Jetz et al. 2019). The need for spatial interpolation may be obvious when working with grids (raster or vector), but it is also present in maps created in non-grid formats, such as with computational geometry that identifies bounding polygons. Interpolation is implicit in the use of bounding polygons around known areas of range or occurrence because an assumption is made that areas between or among areas of known range are also range. Other interpolation approaches could include setting rules about locations relative to known range (e.g. that areas within a specified distance of several occurrence records are within range) and species distribution models (SDMs), which are extremely valuable tools for interpolation with grid formats. Importantly, SDMs often require knowledge of the species’ range when choosing an appropriate modeling extent to avoid extrapolating presence beyond range limits (Guisan et al. 2017), and thus have limited utility for range *delineation* (Graham and Hijmans 2006).

Spatial interpolation methods can also have a temporal component. Data gaps for one period can be filled based on assumptions about a lack of change in range from a previous period for which data were available or for which maps were previously compiled. For example, under an assumption of temporal stationarity (i.e., no change), a spatial unit that is lacking data for a period of interest could be classified as range if it was previously identified as being within the range limits. A similar assumption is used as a basis for combining current and legacy data to produce maps of current distributions under frameworks that do not specify multiple time periods (Jetz et al. 2019).
