## Supplemental Text 2 for "A Framework for Transparent and Repeatable Species Range Maps"

**Map Compilation Details**

Range compilation is programmed in a single Python script that utilizes parallel processing to compile presence and seasonal range maps. The compilation script is written as a collection of custom Python functions that each perform a discrete task:

**Retrieve Compilation Parameters**

Parameters are read from the Parameters Database and set as variables.

**Acquire the GAP Version 1 Data**

Range data are downloaded from ScienceBase with the *sciencebasepy* package (Long et al. 2023) and unzipped to a working directory. This step can be bypassed if no version 1 data is available, useful, or necessary.

**Preparing Species Occurrence Record Geometries**

Although Wildlife Wrangler output databases contain spatial information about records, they do not store the records as spatial objects. The Wildlife Wrangler includes a helper function that generates a shapefile of records as polygons. That function is deployed in the compiler to create geometries for each record, whereby the polygons are record coordinates buffered to a distance equal to the locational uncertainty of the record. However, if a different polygon was specified by the provider for a record, such as a survey block boundary, that shape is used instead. This step can be bypassed if no occurrence records are available, useful, or necessary.

**Create an Output Database**

An SQLite database is created and saved to disk to store compilation input and output. Several tables are created during this step and then populated by subsequent processes.

**Insert Species Occurrence Records**

Occurrence records are read from one or more Wildlife Wrangler output databases and written to a table that includes geometries. Records from unwanted years and months are filtered out.

***Insert Opinion Records***

Records are read from the opinions database and inserted into a table while addressing duplicated, negated, outdated, or conflicting records. Additionally, some opinions for range imply an opinion about presence, and vice versa. For example, an opinion that a spatial unit was within the year-round range of a species also means that the expert believes the species occurred within the unit.

To address those issues, once all opinion records for the species are read into memory, all but the newest records are dropped in cases of duplication for a spatiotemporal unit (unique year and spatial unit pair). Next, all records for a spatiotemporal unit are dropped if conflicting opinions were entered from experts with identical ranks and confidence scores because such records negate each other. In cases where opinions conflict and expert ranks or confidence scores are unequal, the one with the higher expert rank is kept, but if ranks are equal, the record with the highest confidence score is retained. This cleaning process ensures that only one opinion regarding presence and the seasonal ranges is applied to each spatiotemporal unit.

After the opinions records are cleaned, they are adjusted and transferred according to what they imply about presence or seasonal ranges. In the opinion database, each opinion specifically applies to either species presence, summer range, winter range, or year-round range. A column named “status” records whether the opinion is a belief of present or absent for species occurrence or range existence. Under our conceptual basis and definitions of range described above, opinions can transfer under the following logic:

- Range implies presence.
- Absence implies non-range.
- Non-range does not imply absence.
- Year-round range implies summer and winter range.
- Summer *or* winter range does not imply year-round range, but summer *and* winter range implies year-round range.

In some cases, various opinions regarding range and presence within a spatiotemporal unit may align, but the associated expert ranks and confidences may not. To simplify resolving those differences, we combine rank and confidence into a weight value where,

Weight = Expert Rank x (Confidence/10)

In practice, the adjustment (reconciliation and transfer) of opinions is a matter of identifying and addressing certain cases of presence and range opinion value pairs.

1. Presence and range conflict – this is addressed by picking the opinion with the highest weight.
2. Species absence with no range records (NULL) – this is addressed by adding range absence records.
3. Range presence with no record for species – this is addressed by adding species presence records denoting presence.
4. Presence and range agreement on absence but with unequal weights– this is addressed by using the higher weight for both records.

A final subprocess in this step creates the references table by looking up citation codes from the opinions data in an internal GAP database (GAP Wildlife Habitat Relationship Database).

**Document Presence and Range**

Whether presence and range were documented in each spatial unit during each period is assessed by evaluating the number, weights, and locations of occurrence records in relation to each spatial unit. Presence or range are considered documented in spatial units where the sum of record weights from the period is greater than or equal to 10. The sums of record weights and the documented status are saved in columns of the presence and range tables.

The process of summing weights by spatial unit is complicated by issues related to the spatial precision of species occurrence records, which is rarely uniform. Occurrence records are best represented as polygons because they represent events with spatial extents (Chapman and Wieczorek 2020), and the size and shape of record extents are determined by whether the record was recorded as a point or assigned to a sampling unit (e.g. survey block). If recorded as a point, the extent is influenced by the path traveled by the observer, the distance at which the species can be detected under the survey methods, GPS accuracy, and the spatial precision of the coordinates (“nominal coordinate precision”). Represented as polygons, occurrence records can have various spatial relationships with a spatial unit depending upon its location and extent: a single occurrence record could intersect two or more units or be contained within one. When occurrence records are contained within a single unit, there is no uncertainty about which spatial unit the observed organism was located within. However, if the record extent overlaps two or more units, some risk of error exists that is associated with assigning the organism to each overlapping spatial unit (Figure S1). For example, if 40% of a record’s extent were in spatial unit A and 60% in spatial unit B, then assigning the record to spatial unit A carries a 60% risk of misallocating the observation (Figure S1).

**Select the Dominant Opinions**

The opinion with the highest weight is selected for each spatial unit and period. The dominant opinions and their weights are recorded in designated columns.

**Assign Codes**

Presence and range codes are assigned to each spatial unit for each period based on a hierarchy for the data sources that are available from existing columns at this stage in the process (Table S4). Documentation of presence (or range) from observational data of sufficient weight is dominant over all other sources. Opinion is dominant over codes from prior time periods or GAP version 1 data for the first period if the weight of the opinion is sufficient.

Appropriate codes are determined by applying rules in a specific sequence to each spatial unit, for each period. Some steps overwrite the results of a previous step to update the code according to the hierarchy:

1. If a GAP version 1 code exists, use it as a basis for the first period. For presence, the version 1 values “known/extant”, “possibly present”, and “potential for presence” become “suspected present” and version 1 values corresponding to extirpation become “suspected absent”. For seasonal range compilations, if a prior code denoted range for the season, then “suspected range” is assigned to the spatial unit.
2. In the case that a presence code existed for the prior period, reuse all codes for the current period unless the code was “documented present”, in which case assign a code for “suspected present” (Figure 2). In the case of seasonal range, “documented range” becomes “suspected range”, and all other codes are reused for the current period.
3. If an opinion exists for presence and all other sources are NULL, code the spatial unit “suspected present” or “suspected absent” according to the opinion. In the case of range, use “suspected range” and “suspected non-range” instead. If the opinion weight is greater than two, overwrite existing codes to “suspected present”, “suspected absent”, “suspected range”, or “suspected non-range”. If the opinion weight is greater than eight, use codes for “likely” instead of “suspected” (Figure 2).
4. If presence or range were documented occurrence records, then assign the code for “documented presence” or “documented range”, respectively (Figure 2).

The above rules play out in the following ways for presence and range:

- A spatial unit for which there is never any occurrence data or expert opinion will remain coded as it was in GAP version 1.
- If presence is documented in a spatial unit for one period, it will be coded as “suspected present” or “suspected range” in the subsequent periods until expert opinion of absence or non-range is recorded, or presence is documented again.
- If an expert registers their opinion that a species was absent in a unit, but then presence is documented with occurrence data, the expert's opinion will be overridden and the unit will be coded as “documented present” or “documented range”.
- If a spatial unit is coded “suspected present” in one period, but expert opinion indicates absence in the next period, then the unit's code will transition from “documented present” to “suspected” or “likely absent”. Analogous results are produced by the range process: “suspected range” would become “suspected non-range” or “likely non-range”.

**Flag Extralimital Spatial Units**

Spatial units with documented presence from occurrence records of extralimital individuals are found and flagged in a designated column. Spatial units are flagged as extralimital if, 1) documentation only occurred during one period and 2) the centroid of the spatial unit is more than a specified cutoff distance from another spatial unit that is coded as present (or range) during the period being assessed. The cutoff distance is assigned on a per-species, per-task basis in the parameters database to account for variation among taxa. Extralimital records are assessed on a per-period basis because range limits can change over time periods.

**Adjust Codes**

Adjustments to codes are sometimes necessary because extralimital spatial units were identified after the initial code assignment. In addition, some assumptions can be made based upon the available codes from all periods for each spatial unit. A final step adjusts codes considering the temporal context for each period, as well as the extralimital records identified. In cases where presence or range was documented in the prior and subsequent periods to the one being assessed and opinion weight is greater than two, codes are elevated to “likely present” or “likely range” (Figure S2). In cases where presence/range was not documented in the period being assessed, there was no opinion registered, and the spatial unit was flagged as extralimital during a period, the code is set to “suspected absent” or “suspected non-range”.

**Calculate Time Since Last Record**

A table is added to the output database that reports how many weeks passed between the last occurrence record and the date of compilation for each spatial unit. This calculation is not central to GAP’s range mapping objectives, but it is easily attainable once the range data have been compiled.

The calculation requires attribution of occurrence records to spatial units, so the issues surrounding misallocation risk when documenting presence apply to this process as well. Thus, records are again represented as polygons, intersected with spatial units, and the resulting record fragments filtered based upon the allocation error risk from the error tolerance parameter. The table stores the date of the assessment, identification code for the record, record weight, and proportion overlap with the spatial unit, in addition to the age in weeks.

**Make a Simple Results Table**

For convenience and clarity, a simplified table of results is added to the output database that contains only presence and seasonal range columns for each spatial unit.

**List of Tests**

Following map compilation, several automated tests are run to verify that the processes ran correctly.

- In the presence table, the “previously documented” column for the current period should be coded “yes” if presence was documented in a previous period.
- In the presence table, if presence was documented for a period, then past presence should be indicated for all subsequent periods.
- In the presence table, historical weight for a given period should be equal to the sum of recent and historical weight for the previous period.
- In the presence table, presence should only be coded as “documented” when summed occurrence record weights are sufficient for a period.
- In the presence table, “documented presence” columns should be coded “true” for periods with presence code equal set to “1” (confirmed presence).
