## Supplemental Text 3 for "A Framework for Transparent and Repeatable Species Range Maps"

**Fisher Occurrence Record Datasets Provided by GBIF**

Austin J, Viani K, Hammond F, Massa M, Sharp S (2024). Historic Wildlife Roadkill Reports from Vermont, USA (1971-2006). Vermont Center for Ecostudies. Occurrence dataset https://doi.org/10.15468/6a4xjj accessed via GBIF.org on 2025-01-23. http://creativecommons.org/publicdomain/zero/1.0/legalcode.

Bentley A, Krejsa D (2024). KUBI Mammalogy Collection. Version 26.82. University of Kansas Biodiversity Institute. Occurrence dataset https://doi.org/10.15468/a3woj7 accessed via GBIF.org on 2025-01-23. http://creativecommons.org/licenses/by/4.0/legalcode.

Bradley J (2025). UWBM Mammalogy Collection (Arctos). University of Washington Burke Museum. Occurrence dataset https://doi.org/10.15468/qziy3w accessed via GBIF.org on 2025-01-23. http://creativecommons.org/publicdomain/zero/1.0/legalcode.

California Academy of Sciences: CAS Mammalogy (MAM) https://doi.org/10.15468/dhbozg accessed via GBIF.org on 2025-01-23. http://creativecommons.org/publicdomain/zero/1.0/legalcode.

Conroy C (2024). MVZ Mammal Collection (Arctos). Version 35.93. Museum of Vertebrate Zoology. Occurrence dataset https://doi.org/10.15468/uwudf9 accessed via GBIF.org on 2025-01-23. http://creativecommons.org/publicdomain/zero/1.0/legalcode.

Cook J (2024). MSB Mammal Collection (Arctos). Version 35.94. Museum of Southwestern Biology. Occurrence dataset https://doi.org/10.15468/oirgxw accessed via GBIF.org on 2025-01-23. http://creativecommons.org/publicdomain/zero/1.0/legalcode.

Galbreath K (2024). Northern Michigan University (NMU) Mammal Specimens (Arctos). Version 1.84. Northern Michigan University. Occurrence dataset https://doi.org/10.15468/zazfgp accessed via GBIF.org on 2025-01-23. http://creativecommons.org/publicdomain/zero/1.0/legalcode.

Gall L (2025). Vertebrate Zoology Division - Mammalogy, Yale Peabody Museum. Yale University Peabody Museum. Occurrence dataset https://doi.org/10.15468/4mm6uc accessed via GBIF.org on 2025-01-23. <http://creativecommons.org/publicdomain/zero/1.0/legalcode>. http://creativecommons.org/publicdomain/zero/1.0/legalcode.

Garretson A, Napoli M, Feldsine N, Long E, Huth P, Smiley D, Forester A, Pierce E, Smiley S, Thompson J (2022). Mohonk Preserve Historical Observational Biodiversity Data. Mohonk Preserve. Occurrence dataset https://doi.org/10.15468/tckm2a accessed via GBIF.org on 2025-01-23. http://creativecommons.org/publicdomain/zero/1.0/legalcode.

Harvard University M, Morris P J (2025). Museum of Comparative Zoology, Harvard University. Version 162.454. Museum of Comparative Zoology, Harvard University. Occurrence dataset https://doi.org/10.15468/p5rupv accessed via GBIF.org on 2025-01-23. http://creativecommons.org/licenses/by-nc/4.0/legalcode.

Kuprewicz E (2020). UConn Mammals. Version 3.3. University of Connecticut. Occurrence dataset https://doi.org/10.15468/dbs8w7 accessed via GBIF.org on 2025-01-23. http://creativecommons.org/publicdomain/zero/1.0/legalcode.

iNaturalist contributors, iNaturalist (2025). iNaturalist Research-grade Observations. iNaturalist.org. Occurrence dataset https://doi.org/10.15468/ab3s5x accessed via GBIF.org on 2025-01-23. <http://creativecommons.org/licenses/by-nc/4.0/legalcode>.

Motz G (2025). Vertebrate Zoology Division - Mammalogy, Yale Peabody Museum. Yale University Peabody Museum. Occurrence dataset https://doi.org/10.15468/4mm6uc accessed via GBIF.org on 2025-01-23. http://creativecommons.org/publicdomain/zero/1.0/legalcode.

UMMZ Mammals Data Group, LSA IT A (2025). University of Michigan Museum of Zoology, Division of Mammals. Version 8.81. University of Michigan Museum of Zoology. Occurrence dataset https://doi.org/10.15468/dx3rcj accessed via GBIF.org on 2025-01-23. http://creativecommons.org/licenses/by-nc/4.0/legalcode.

University of Wisconsin – Stevens Point (2025). University of Wisconsin Stevens Point Mammals. Occurrence dataset https://doi.org/10.15468/7mp2c4 accessed via GBIF.org on 2025-01-23. http://creativecommons.org/licenses/by/4.0/legalcode.

Whitehouse R, Gerhard D (2021). NYSM Mammals. Version 16.2. New York State Museum (NYSM). Occurrence dataset https://doi.org/10.15468/awwifu accessed via GBIF.org on 2025-01-23. <http://creativecommons.org/publicdomain/zero/1.0/legalcode>.
