## Supplemental Text 4 for "A Framework for Transparent and Repeatable Species Range Maps"

**Sources for Secondary Expert Opinions about Fisher Range and Presence**

Coltrane J, Inman R. 2021. Fisher occupancy twenty-five years after translocation in the Rocky Mountains of Montana. Northwestern Naturalist 102:43–54. https://doi.org/10.1898/1051-1733-102.1.43.

*Range absence was supported for 28 spatial units because the authors performed systematic trapping in the Cabinet Mountains of Montana and only captured fishers that were related to translocated individuals, not an established population.*

Green DS, Facka AN, Smith KP, Matthews SM, Powell RA. 2022. Evaluating the efficacy of reintroducing fishers (Pekania pennanti) to a landscape managed for timber production. Forest Ecology and Management 511: 120089. https://doi.org/10.1016/j.foreco.2022.120089.

*Range presence was supported for 612 spatial units because the authors detailed the successful reintroduction of fishers.*

Green RE. 2017. Reproductive ecology of the fisher (Pekania pennanti) in the southern Sierra Nevada: An assessment of reproductive parameters and forest habitat used by denning females. Doctoral dissertation. Davis, California: University of California Davis.

*Range presence was supported for 270 spatial units because the authors sampled individuals from an extant population.*

Happe PJ, Jenkins KJ, Mccaffery RM, Lewis JC, Pilgrim KL, Schwartz MK. 2020. Occupancy Patterns in a Reintroduced Fisher Population during Reestablishment. Journal of Wildlife Management 84: 344-358. https://doi.org/10.1002/jwmg.21788.

*Range presence was supported for 572 spatial units because this study suggested that the reintroduced population is now established. Presence was also supported for 94 spatial units where fishers were detected.*

Keinath DA, Andersen MD, Beauvais GP. 2010. Range maps for Wyoming species of greatest conservation need. Report of Wyoming Natural Diversity Database to Wyoming Game and Fish Department, Cheyenne, Wyoming and the U.S. Geological Survey, Fort Collins, Colorado. Available: https://www.uwyo.edu/wyndd/_files/docs/reports/wynddreports/u10kei01wyus.pdf(April 2024).

*The dataset coded the species as present but “irregular” in 1,600 spatial units, which supported their inclusion in the presence map, but not the range map.*

Krohner JM, Lukacs PM, Inman R, Sauder JD, Gude JA, Mosby C, Coltrane JA, Mowry RA, Millspaugh JJ. 2022. Finding fishers: Determining fisher occupancy in the Northern Rocky Mountains. Journal of Wildlife Management 86:e22162. https://doi.org/10.1002/jwmg.22162.

*The authors modeled fisher occupancy of the northern Rocky Mountains and their predictions supported the removal of 2,002 spatial units and addition of 318 units to the range map. They reported locations of fisher detections that supported the addition of 152 units to the presence map.*

Lewis JC, Powell RA, Zielinski WJ. 2012. Carnivore translocations and conservation: Insights from population models and field data for Fishers (Martes pennanti). PLoS ONE 7: e32726. doi:10.1371/journal.pone.0032726.

*The authors provided a map of historic and current (as of 2012) range that closely resembles the map that we compiled.*

Lewis JC, Ransom JI, Chestnut T, Werntz DO, Black S, Whiteside D, Postigo JL, Moehrenschlager A. 2022. Cascades fisher reintroduction project: Final project report. Fort Collins, CO: NPS. Natural Resource Report NPS/PWR/NRR—2022/2418. <https://doi.org/10.36967/2293605>.

*Range presence was supported for 912 spatial units because the authors provided a map of fisher detections.*

Lucid MK, Rankin A, Sullivan J, Robinson L, Ehlers S, Cushman S. 2019. A carnivore's oasis? An isolated fisher (Pekania pennanti) population provides insight on persistence of a metapopulation. Conservation Genetics 20:585-596. https://doi.org/10.1007/s10592-019-01160-w.

*The authors conducted a thorough genetic study of fisher populations that supported the presence of range within 860 spatial units.*

Moncrief ND, Fies ML. 2015. Report of first specimens of Pekania pennanti (Fisher) from Virginia. Northeastern Naturalist 22(4): 31-34. https://doi.org/10.1656/045.022.0417.

*The authors described fisher records that indicated presence in 45 spatial units, but they determined that the individuals were not within the species’ range.*

National Park Service. 2021. First wild fishers born in the North Cascades. News Release. Available: https://www.nps.gov/noca/learn/news/first-wild-fishers-born-in-the-north-cascades.htm. (March 2023).

Parsons MA, Lewis JC, Puali JN, Chestnut T, Ransom JI, Werntz DO, Prugh LR. 2020. Prey of reintroduced fishers and their habitat relationships in the Cascades Range, Washington. Forest Ecology and Management 460:117888. https://doi.org/10.1016/j.foreco.2020.117888.

*The authors provided a map of sample sites from an established population that indicated that 196 spatial units contained a portion of the range.*

Pauli JN, Manlick PJ, Tucker JM, Smith GB, Jensen PG, Fisher JT. 2022. Competitive overlap between martens Martes americana and Martes caurina and fishers Pekania pennanti: A rangewide perspective and synthesis. Mammal Review 52: 392-409. https://doi.org/10.1111/mam.12284.

*The authors provided a small, coarse range map that suggested absence of range in 3,685 spatial units and presence of range in 7,884 spatial units.*

Powell RA. 1981. Martes pennanti. Mammalian Species 156:1-6.

*This species account included a map that identified areas from which the species was extirpated that implied absence in 365 spatial units.*

Sweitzer RA, Popescu VD, Barret RH, Purcell KL, Thompson CM. 2021. Reproduction, abundance, and population growth for a fisher (Pekania pennanti) population in the Sierra National Forest, California. Journal of Mammalogy 96:772-790. https://doi.org/10.1093/jmammal/gyv083.

*Range presence in 248 spatial units was supported by maps provided by the authors.*

Tucker JM. 2013. Assessing changes in connectivity and abundance through time for fisher in the southern sierra nevada. PhD dissertation. Missoula, Montana: University of Montana.

*This publication supports belief in the absence of fishers from a section of the Sierra Nevadas that spanned 642 spatial units.*

Waller JS. 2018. Status of fishers in Glacier National Park, Montana. Northwestern Naturalist 99(1):1-8. https://doi.org/10.1898/NWN17-07.1.

*Extensive surveying and review of occurrence records supported belief that 909 spatial units were in an area where fishers occurred, but that was not part of the species’ range. 371 spatial units that included an area where fishers were reestablished were also identified by the authors.*
