## Supplemental Figure 1 for "A Framework for Transparent and Repeatable Species Range Maps"

### Slide 1
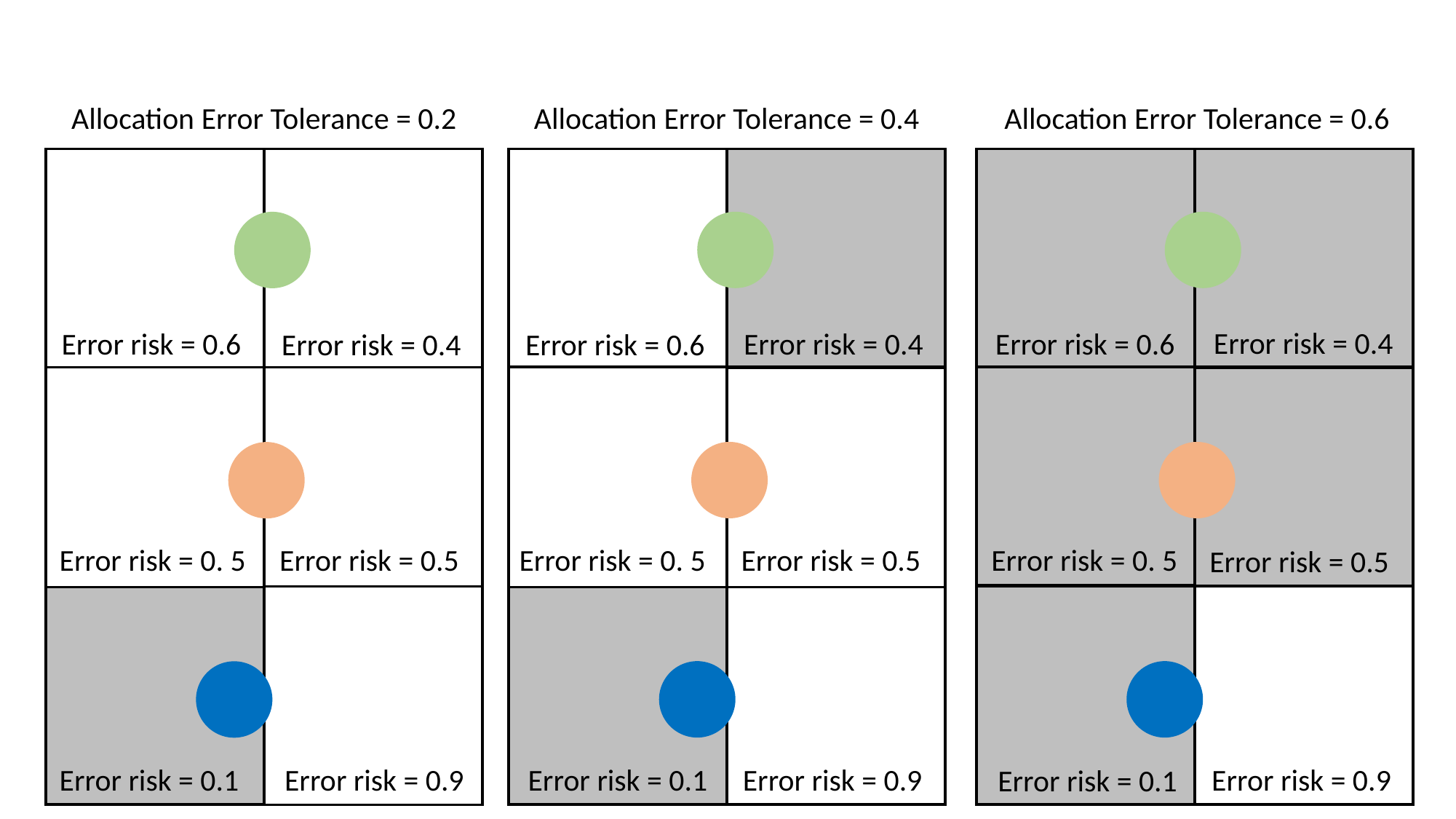

Allocation Error Tolerance = 0.6
Error risk = 0.4
Error risk = 0.6
Error risk = 0. 5
Error risk = 0.5
Error risk = 0.9
Error risk = 0.1
Allocation Error Tolerance = 0.2
Error risk = 0.6
Error risk = 0.4
Error risk = 0. 5
Error risk = 0.5
Error risk = 0.9
Error risk = 0.1
Allocation Error Tolerance = 0.4
Error risk = 0.4
Error risk = 0.6
Error risk = 0. 5
Error risk = 0.5
Error risk = 0.9
Error risk = 0.1
