## Supplementary figures and images for "A Framework for Transparent and Repeatable Species Range Maps"

### Supplemental Figure 2

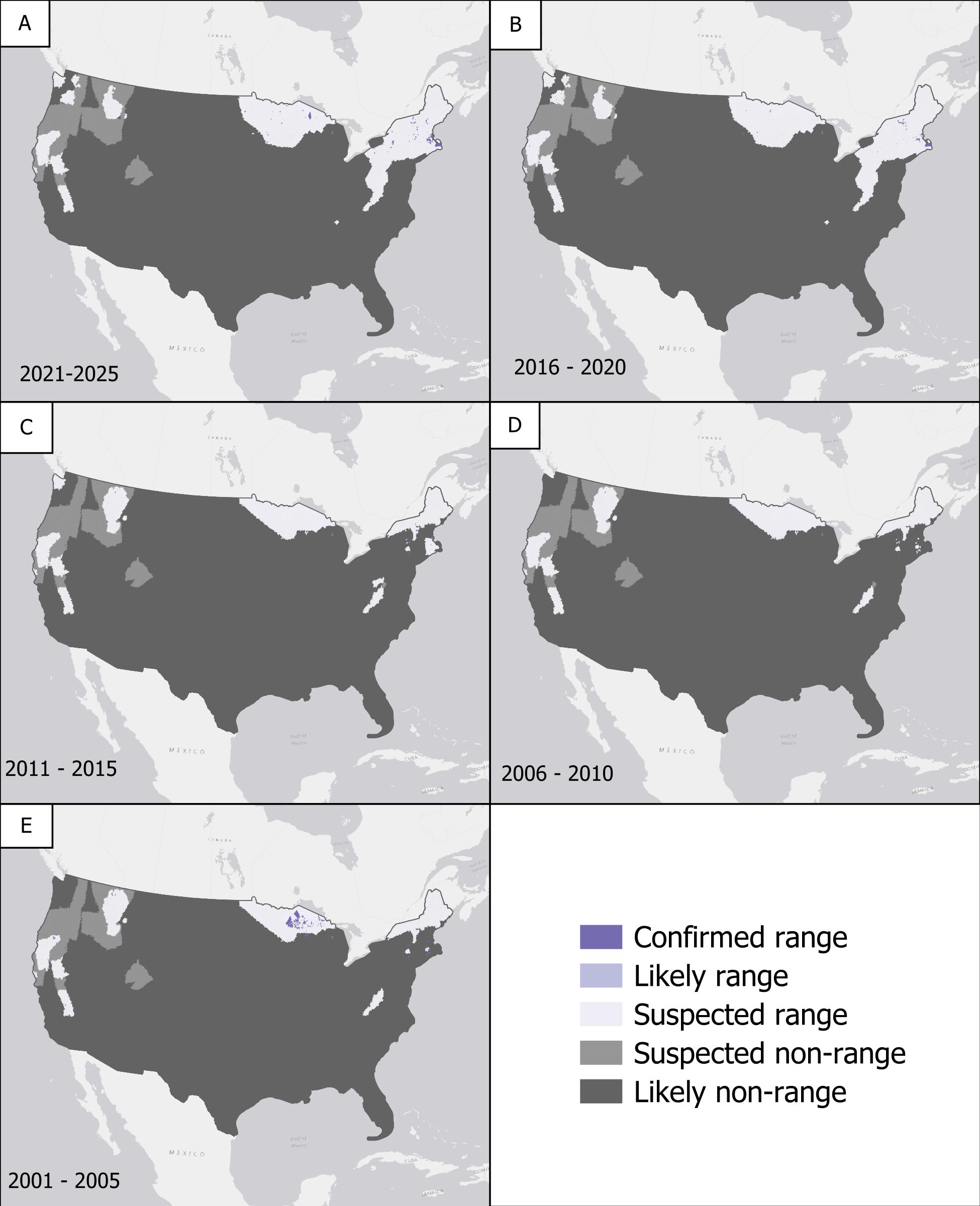

### Supplemental Figure 3

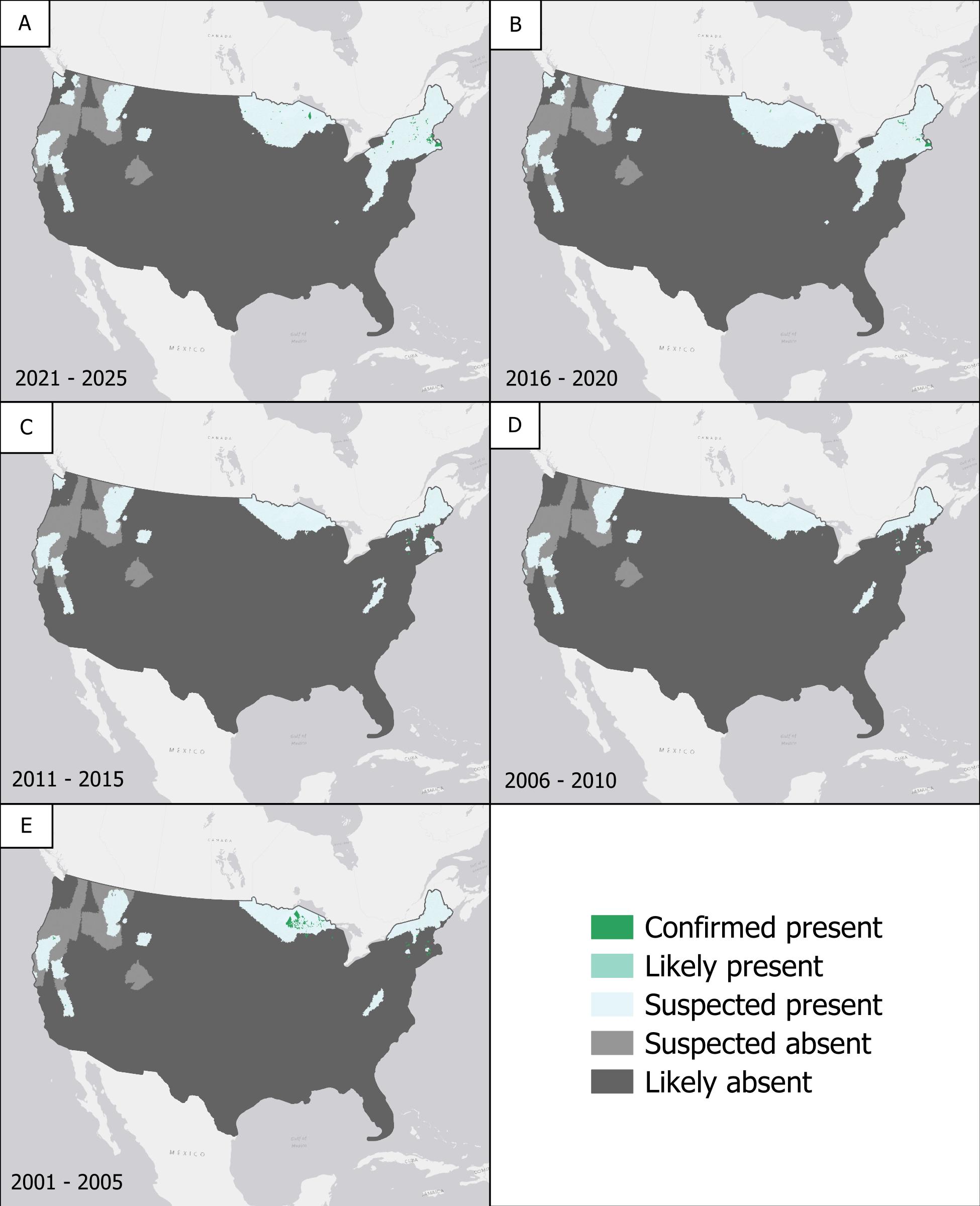

### Supplemental Figure 4

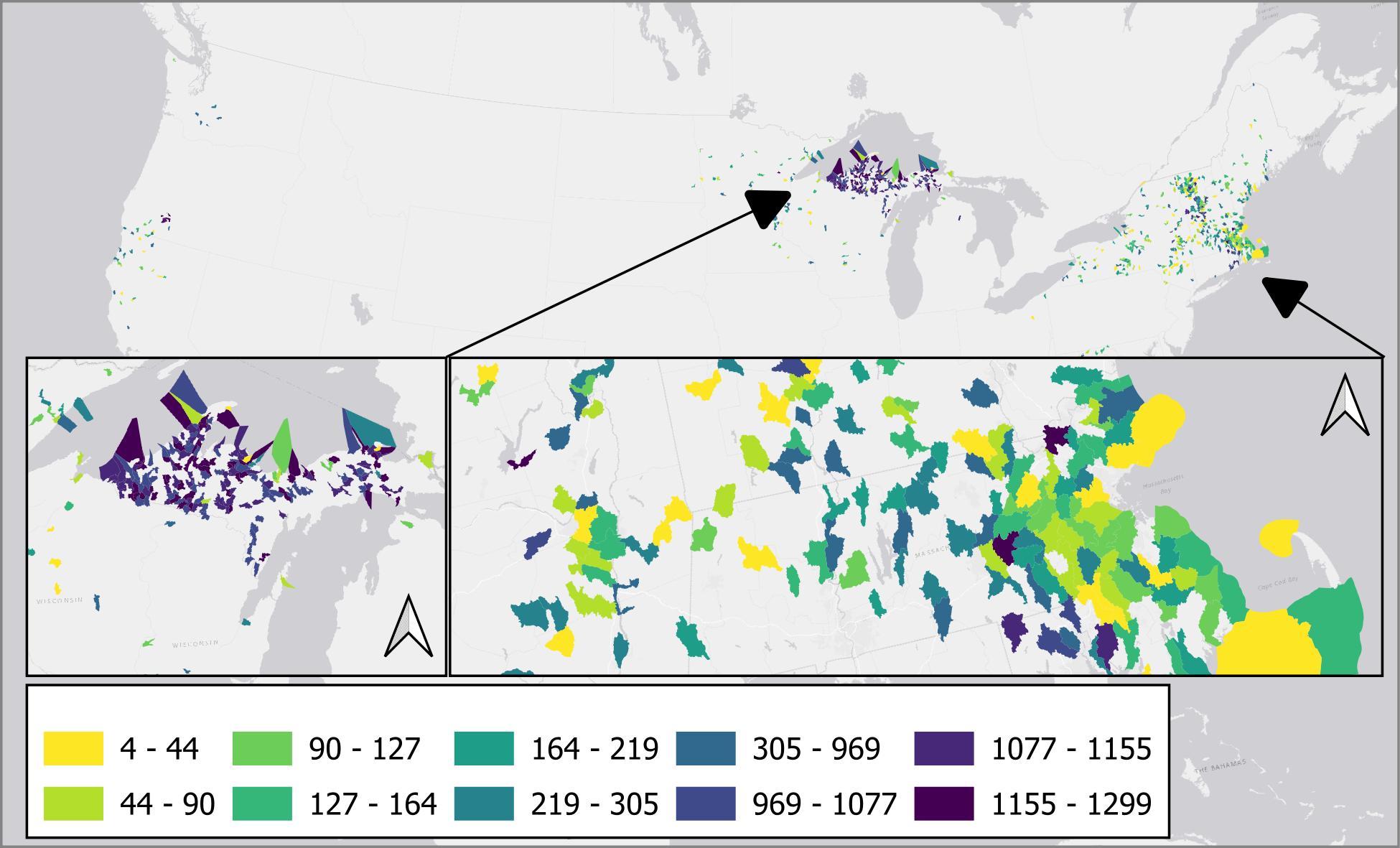

### Supplemental Figure 5

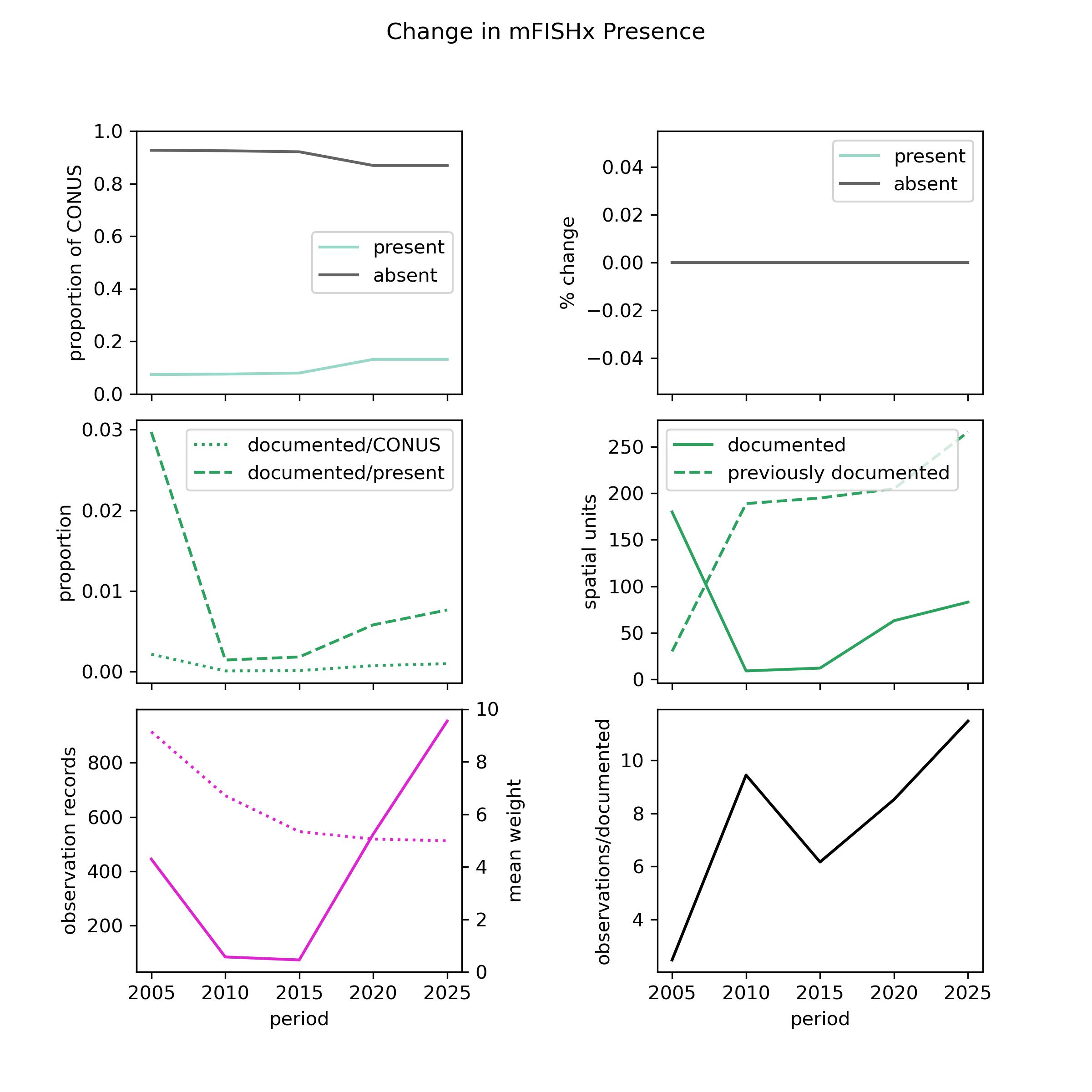

### Supplemental Figure 6

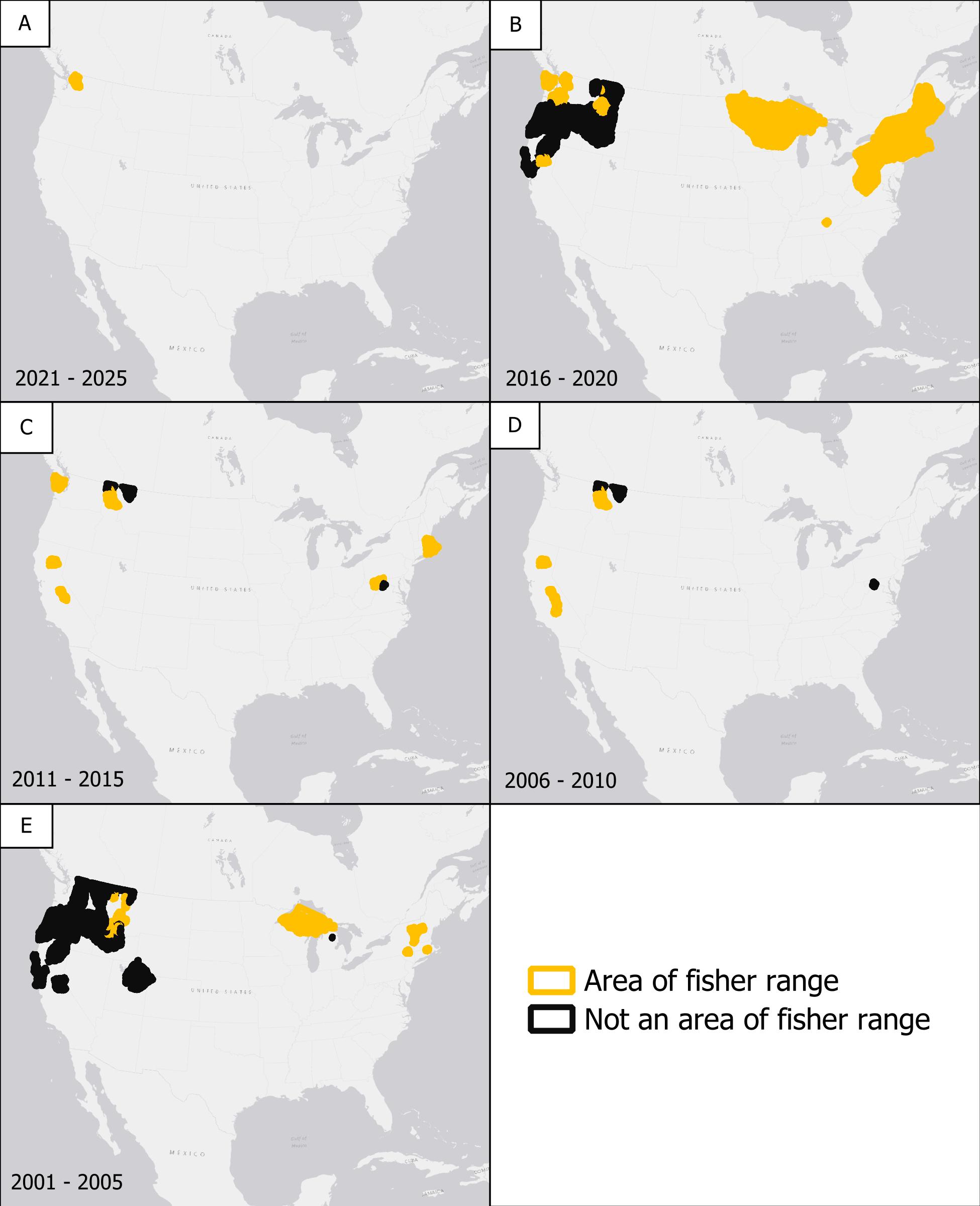

### Supplemental Figure 7

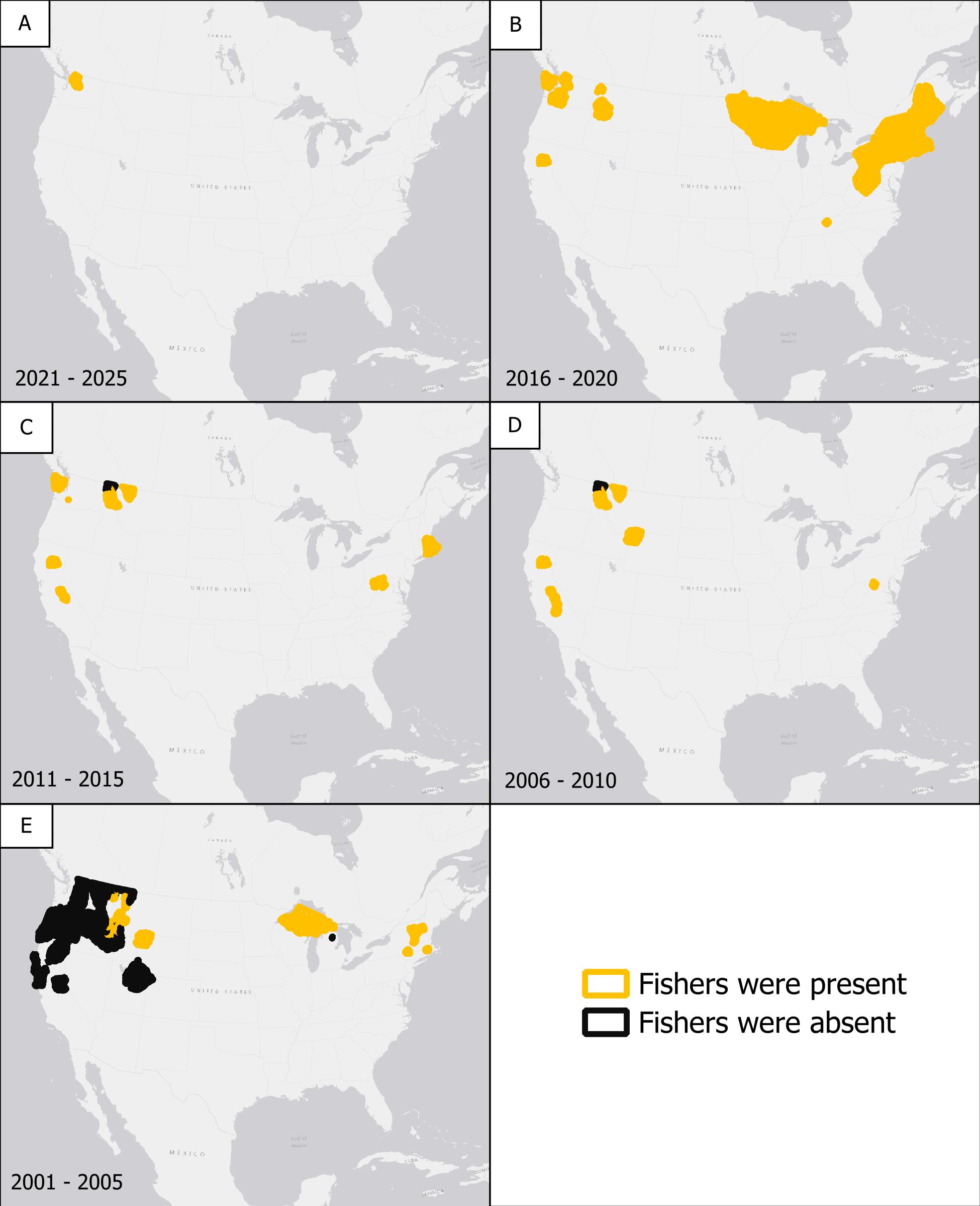
