## Supplemental Figure 8 for "A Framework for Transparent and Repeatable Species Range Maps"

### Slide 1
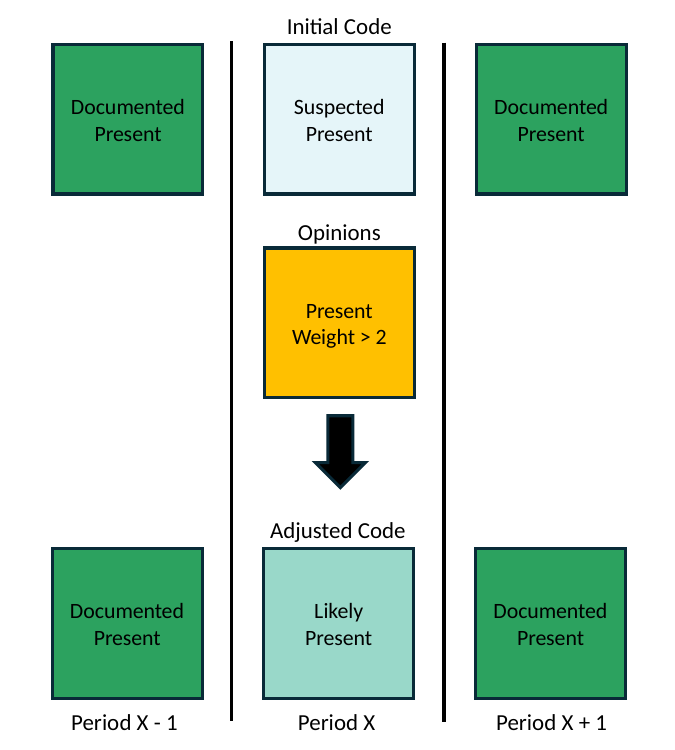

Initial Code
Documented Present
Suspected Present
Documented Present
Opinions
Present
Weight > 2
Adjusted Code
Documented Present
Likely Present
Documented Present
Period X - 1
Period X
Period X + 1
