## Supplementary material for "A Framework for Transparent and Repeatable Species Range Maps": Occurrence Record Queries: File S1.html

mFISHx0\_Observations


### Fisher Records from GBIF¶

```
Notebook run 2025-01-23 13:40:43.281207
Results were saved in C:/Workspaces/RangeMaps/fisher/mFISHx0_Observations.sqlite
```

#### Taxon Concept¶

```
{'EBIRD_ID': 'None',
 'GBIF_ID': 8631083,
 'ID': 'mfishx0',
 'TAXON_EOO': None,
 'common_name': 'fisher',
 'detection_distance_m': 50,
 'scientific_name': 'Pekania pennanti'}
```

#### Filter Set¶

```
Filter set name: Fisher0
```

###### GBIF Request Method¶

```
Request a Darwin Core Archive? True
```

###### Date Limits¶

Notes: The taxa concept was revised between 2012 and 2014, but older records that were attributed to Martes pennanti may have been adjusted, so pre-2012 records were included.

```
Years: 2000,2024
Months: 1,12
```

###### Country¶

Notes:

```
Country: US
```

###### Bounding Box¶

Notes:

```
Latitude range: None
Longitude range: None
```

###### Area of Interest¶

Notes:

```
None
```

###### Taxon EOO¶

Notes:

```
Use taxon extent of occurrence? False
```

###### Geoissue¶

Notes:

```
Records with geoissues OK? False
```

###### Collections¶

Notes:

```
Omit: None
```

###### Institutions¶

Notes:

```
Omit: None
```

###### Datasets¶

Notes:

```
Omit: None
```

###### Coordinate Uncertainty¶

Notes:

```
Coordinate uncertainty required? False
Default coordinate uncertainty to use: None
Maximum allowable coordinate uncertainty: 100000
```

###### Bases¶

Notes:

```
Omit: None
```

###### Sampling Protocols¶

Notes:

```
Omit: None
```

###### Issues¶

Notes: Invalid dates could create errors in the removal of duplicates and/or assigning records to time periods. Taxon matching needs to be correct so that other species' records are not included. Invalid type statuses could allow the inclusion of fossils.

```
Omit: ['TAXON_MATCH_HIGHERRANK', 'TYPE_STATUS_INVALID', 'RECORDED_DATE_INVALID']
```

###### Duplicates¶

Notes:

```
Allow duplicates? False
```

###### Filter Set Summary¶

```
{'bases_omit': None,
 'collection_codes_omit': None,
 'country': 'US',
 'datasets_omit': None,
 'default_coordUncertainty': None,
 'duplicate_coord_date_OK': False,
 'geoissue': False,
 'get_dwca': True,
 'has_coordinate_uncertainty': False,
 'institutions_omit': None,
 'issues_omit': ['TAXON_MATCH_HIGHERRANK',
                 'TYPE_STATUS_INVALID',
                 'RECORDED_DATE_INVALID'],
 'lat_range': None,
 'lon_range': None,
 'max_coordinate_uncertainty': 100000,
 'months_range': '1,12',
 'name': 'Fisher0',
 'query_polygon': None,
 'sampling_protocols_omit': None,
 'use_taxon_geometry': False,
 'years_range': '2000,2024'}
```

#### Processing¶

```
Prepared filter set and sorted out geometry constraints: 0:00:00
2444 records available
```

```
INFO:Your download key is 0003098-250121130708018
```

```
Waiting for the Darwin Core Archive.....
```

```
INFO:Download file size: 717499 bytes
INFO:On disk at C:/Workspaces/RangeMaps/fisher//0003098-250121130708018.zip
```

```
Wait time for DWcA creation: 0:01:18.671873
Wait time for DWcA download: 0:00:02.187363
Wait time for reading the DwCA: 0:00:00.569527
Stored GBIF Download DOI etc.: 0:00:00
Summarized fields returned: 0:00:00.078999
Prepared GBIF records for processing: 0:00:00.095019
```

```
Prepared data frames for processing: 0:00:00.024582
Summarized values acquired: 0:00:00.078251
An unsupported date format has been found:  2001-01
An unsupported date format has been found:  2001-12
An unsupported date format has been found:  2005-10
An unsupported date format has been found:  2001-02
An unsupported date format has been found:  2001-02
An unsupported date format has been found:  2016-08
An unsupported date range has been found:  ['2014-12-01', '2014-12-31']
An unsupported date range has been found:  ['2018-12-01', '2018-12-31']
An unsupported date range has been found:  ['2016-02-01', '2016-02-29']
An unsupported date range has been found:  ['2019-01-01', '2019-01-31']
An unsupported date range has been found:  ['2018-01-01', '2018-01-31']
An unsupported date range has been found:  ['2013-12-01', '2013-12-31']
Parsed event dates: 0:00:00.035050
Calculated nominal coordinate precision: 0:00:00.032630
Number of georeferenced GBIF records: 1460
Approximating coordinate uncertanties for GBIF records
Prepared records and calculated radii:0:00:00.071514
Performed filtering: 0:00:00.032987
138 duplicate records dropped: 0:00:00.388513
Saved summary of filtering results: 0:00:00.101067
```

```
2287 records were saved in the output database
```

#### Results of the Filtering¶

###### Sources¶

```
                     institutionID collectionCode                     datasetName  acquired  removed  retained
0                              MVZ           Mamm                             nan         2        0         2
1                             UWBM           Mamm                             nan        21        2        19
2   b4640710-8e03-11d8-b956-b8a...           Mamm                             nan        30        0        30
3   http://grbio.org/cool/iakn-...            KUM  University of Kansas Biodiv...       604       89       515
4   http://grbio.org/cool/jrf7-...        Mammals                             nan         1        0         1
5   http://grscicoll.org/instit...            MAM                             nan         1        0         1
6                              nan        Mammals                             nan         2        1         1
7                              nan   Observations  iNaturalist research-grade ...      1584       29      1555
8                              nan             VZ                             nan         4        2         2
9                              nan             ZM                             nan        55       18        37
10                             nan            nan                             nan       132        8       124
```

###### Bases¶

```
                value  acquired  removed  retained
0   HUMAN_OBSERVATION      1716       37      1679
1  PRESERVED_SPECIMEN       728      120       608
```

###### Protocols¶

```
                value  acquired  removed  retained
0        Dead on Road         1        0         1
1           Road kill         1        0         1
2                 nan      2429      157      2272
3            roadkill         1        0         1
4  salvage (roadkill)         9        0         9
5                shot         1        0         1
6             trapped         2        0         2
```

###### Issues¶

```
                                                                                        value  acquired  removed  retained
0           CONTINENT_DERIVED_FROM_COORDINATES;INSTITUTION_MATCH_FUZZY;COLLECTION_MATCH_FUZZY        18        1        17
1                             CONTINENT_DERIVED_FROM_COORDINATES;TAXON_MATCH_TAXON_ID_IGNORED       115        9       106
2   CONTINENT_DERIVED_FROM_COORDINATES;TAXON_MATCH_TAXON_ID_IGNORED;INSTITUTION_MATCH_FUZZ...         2        1         1
3   COORDINATE_REPROJECTED;CONTINENT_DERIVED_FROM_COORDINATES;INSTITUTION_MATCH_FUZZY;COLL...         2        1         1
4   COORDINATE_ROUNDED;CONTINENT_DERIVED_FROM_COORDINATES;INSTITUTION_MATCH_FUZZY;COLLECTI...         1        0         1
5          COORDINATE_ROUNDED;CONTINENT_DERIVED_FROM_COORDINATES;TAXON_MATCH_TAXON_ID_IGNORED      1469       20      1449
6   COORDINATE_ROUNDED;COORDINATE_REPROJECTED;CONTINENT_DERIVED_FROM_COORDINATES;INSTITUTI...         1        0         1
7                      COORDINATE_ROUNDED;GEODETIC_DATUM_INVALID;GEODETIC_DATUM_ASSUMED_WGS84         1        0         1
8                  COORDINATE_ROUNDED;INSTITUTION_MATCH_FUZZY;INSTITUTION_COLLECTION_MISMATCH         3        0         3
9   COORDINATE_ROUNDED;OCCURRENCE_STATUS_INFERRED_FROM_INDIVIDUAL_COUNT;INSTITUTION_MATCH_...         4        2         2
10  GEODETIC_DATUM_INVALID;GEODETIC_DATUM_ASSUMED_WGS84;CONTINENT_DERIVED_FROM_COORDINATES...         7        6         1
11                                             INSTITUTION_MATCH_FUZZY;COLLECTION_MATCH_FUZZY        30        0        30
12                                    INSTITUTION_MATCH_FUZZY;INSTITUTION_COLLECTION_MISMATCH        52       18        34
13                                           OCCURRENCE_STATUS_INFERRED_FROM_INDIVIDUAL_COUNT       106        7        99
14                                                                                        nan       631       90       541
```

Unique issues that were associated with records:

```
COORDINATE_REPROJECTED
OCCURRENCE_STATUS_INFERRED_FROM_INDIVIDUAL_COUNT
COLLECTION_MATCH_FUZZY
TAXON_MATCH_TAXON_ID_IGNORED
CONTINENT_DERIVED_FROM_COORDINATES
COORDINATE_ROUNDED
INSTITUTION_COLLECTION_MISMATCH
GEODETIC_DATUM_INVALID
INSTITUTION_MATCH_FUZZY
GEODETIC_DATUM_ASSUMED_WGS84
nan
```

###### Establishment Means¶

```
    value  acquired  removed  retained
0     nan      2442      156      2286
1  native         2        1         1
```

###### Identification Qualifiers¶

```
No identification qualifiers were reported.
```

#### Descriptions of Retained Records¶

###### Locations¶

###### Years Represented¶

Out[34]:

```
Text(0.5, 1.0, 'Occurrences per Year')
```

###### Months Represented¶

Out[35]:

```
Text(0.5, 1.0, 'Occurrences per Month')
```

###### Distribution of Coordinate Uncertainty Values for Retained Records¶

###### Distribution of Point-radius Values for Retained Records¶

Out[38]:

```
Text(0.5, 1.0, 'Compiled Point-radius Buffer Lengths')
```

###### Distribution of Nominal Coordinate Precisions for Retained Records¶

Out[39]:

```
Text(0.5, 1.0, 'Nominal Precisions of Coordinates')
```

###### Remarks¶

```
General remarks:
<NA>
```

```
Event remarks:
<NA>
Former verification status: verified by curator.
native; missing all feet; neck circ=180mm
native; "Phillips kit male"; ear punch dry & refilled JAN05
native; "Phillip's Cr. F - 21"
native; "Adult Male M-24"
native; "Curr Creek F-22"
native; "Quarry F-20"; ear punch dry & refilled JAN 2005
native; "Rustler Peak F-18"
native; "New Guy M-23"
native; "Middlefork F-19"; ear punch dry & refilled JAN05
native; "Foster Cr male - 02"; flat skin
native; "Union Creek Female-11"; necropsied by collector
Likely raptor kill but unconfirmed
```

```
Occurrence remarks:
More than 20 remarks, consult the occurrence database.
```

```
Location remarks:
More than 20 remarks, consult the occurrence database.
```

```
Identified remarks:
<NA>
```

```
Georeference remarks:
<NA>
provisional georeference to ST CO PL: Connecticut New Haven Guilford, 6 Mar 2015, LFG
radius inferred 2024-01-04 from coordinates
Accurate to general area.
[UNDER REVIEW; GEOLOCATE_SCORE: 36; GEOLOCATE_PRECISION: Low; GEOLOCATE_NUMRESULTS: 1]
marker placed on Angell Rd. Error polygon includes entire length of road
Marker placed on Rte 116 in Smithfield, RI
marker placed on rte 135, 0.10 miles south of rte 9 junction
Based off location of Singletary Lane, Sudbury, MA
Based off location of Woburn Street, Andover MA
marker placed on Eastern Ave, north of Pond Rd
Based off address of 272 Lowell St, Andover, MA
Based off location of Lincoln Circle, Andover, MA
```

###### Attributes Returned for GBIF Records¶

This count was made before filters were applied

```
                                 attribute  included(n)  populated(n)
Field                                                                
148                      acceptedNameUsage         2444             0
141                    acceptedNameUsageID         2444             0
201                 acceptedScientificName         2444          2444
191                       acceptedTaxonKey         2444          2444
0                             accessRights         2444           694
41                   associatedOccurrences         2444             1
50                     associatedOrganisms         2444             0
42                    associatedReferences         2444             1
43                     associatedSequences         2444             0
44                          associatedTaxa         2444             0
16                           basisOfRecord         2444          2444
128                                    bed         2444             0
32                                behavior         2444            70
1                    bibliographicCitation         2444            60
31                                   caste         2444             0
21                           catalogNumber         2444          2338
157                                  class         2444          2444
194                               classKey         2444          2444
13                          collectionCode         2444          2312
10                            collectionID         2444            33
79                               continent         2444          2444
99                     coordinatePrecision         2444            30
98           coordinateUncertaintyInMeters         2444          1466
83                             countryCode         2444          2444
85                                  county         2444           731
170                        cultivarEpithet         2444             0
18                     dataGeneralizations         2444           106
11                               datasetID         2444             4
178                             datasetKey         2444          2444
14                             datasetName         2444          2188
135                         dateIdentified         2444          1610
67                                     day         2444          2432
96                         decimalLatitude         2444          2444
97                        decimalLongitude         2444          2444
35                   degreeOfEstablishment         2444             0
183                                  depth         2444             0
184                          depthAccuracy         2444             0
40                             disposition         2444            34
185           distanceFromCentroidInMeters         2444             0
19                       dynamicProperties         2444            64
120               earliestAgeOrLowestStage         2444             0
112            earliestEonOrLowestEonothem         2444             0
118            earliestEpochOrLowestSeries         2444             0
114             earliestEraOrLowestErathem         2444             0
116           earliestPeriodOrLowestSystem         2444             0
181                              elevation         2444            18
182                      elevationAccuracy         2444            18
64                            endDayOfYear         2444          2438
34                      establishmentMeans         2444             2
61                               eventDate         2444          2444
57                                 eventID         2444             0
75                            eventRemarks         2444            14
62                               eventTime         2444          1529
59                               eventType         2444             0
160                                 family         2444          2444
196                              familyKey         2444          2444
74                              fieldNotes         2444             0
60                             fieldNumber         2444            11
104                           footprintSRS         2444             3
105                    footprintSpatialFit         2444             0
103                           footprintWKT         2444            33
126                              formation         2444             0
211                             gbifRegion         2444          2444
165                            genericName         2444          2444
164                                  genus         2444          2444
197                               genusKey         2444          2444
222                          geodeticDatum         2444          2444
111                    geologicalContextID         2444             0
108                   georeferenceProtocol         2444           147
110                    georeferenceRemarks         2444            14
109                    georeferenceSources         2444           681
37          georeferenceVerificationStatus         2444            87
106                        georeferencedBy         2444           693
107                      georeferencedDate         2444           649
125                                  group         2444             0
69                                 habitat         2444             1
188                          hasCoordinate         2444          2444
189                    hasGeospatialIssues         2444          2444
154                   higherClassification         2444           124
78                         higherGeography         2444           122
77                       higherGeographyID         2444            28
123            highestBiostratigraphicZone         2444             0
129                       identificationID         2444          1579
131                identificationQualifier         2444            30
136               identificationReferences         2444             0
138                  identificationRemarks         2444            93
137       identificationVerificationStatus         2444             0
133                           identifiedBy         2444          1640
134                         identifiedByID         2444            35
25                         individualCount         2444           221
17                     informationWithheld         2444           296
167                    infragenericEpithet         2444             0
169                   infraspecificEpithet         2444           637
12                         institutionCode         2444          2312
9                            institutionID         2444           666
210                            isSequenced         2444          2444
82                                  island         2444             0
81                             islandGroup         2444             0
186                                  issue         2444          1813
221                    iucnRedListCategory         2444          1807
155                                kingdom         2444          2444
192                             kingdomKey         2444          2444
2                                 language         2444            91
206                            lastCrawled         2444          2444
180                        lastInterpreted         2444          2444
205                             lastParsed         2444          2444
121                latestAgeOrHighestStage         2444             0
113             latestEonOrHighestEonothem         2444             0
119             latestEpochOrHighestSeries         2444             0
115              latestEraOrHighestErathem         2444             0
117            latestPeriodOrHighestSystem         2444             0
213                              level0Gid         2444          2436
214                             level0Name         2444          2436
215                              level1Gid         2444          2436
216                             level1Name         2444          2436
217                              level2Gid         2444          2436
218                             level2Name         2444          2436
219                              level3Gid         2444             0
220                             level3Name         2444             0
3                                  license         2444          2444
29                               lifeStage         2444           280
124                lithostratigraphicTerms         2444             0
87                                locality         2444           834
94                     locationAccordingTo         2444            30
76                              locationID         2444             0
95                         locationRemarks         2444           619
122             lowestBiostratigraphicZone         2444             0
53                        materialEntityID         2444             0
54                   materialEntityRemarks         2444             0
56                        materialSampleID         2444             0
93     maximumDistanceAboveSurfaceInMeters         2444             0
187                              mediaType         2444          1488
127                                 member         2444             0
92     minimumDistanceAboveSurfaceInMeters         2444             0
4                                 modified         2444          2312
66                                   month         2444          2444
86                            municipality         2444            30
151                        nameAccordingTo         2444             0
144                      nameAccordingToID         2444             0
152                        namePublishedIn         2444             0
145                      namePublishedInID         2444             0
153                    namePublishedInYear         2444             0
174                      nomenclaturalCode         2444           114
176                    nomenclaturalStatus         2444             0
20                            occurrenceID         2444          2444
46                       occurrenceRemarks         2444          1245
38                        occurrenceStatus         2444          2444
158                                  order         2444          2444
195                               orderKey         2444          2444
47                              organismID         2444            30
48                            organismName         2444             0
26                        organismQuantity         2444             0
27                    organismQuantityType         2444             0
52                         organismRemarks         2444             0
49                           organismScope         2444             0
150                      originalNameUsage         2444             0
143                    originalNameUsageID         2444             0
45                     otherCatalogNumbers         2444            59
15                    ownerInstitutionCode         2444            34
58                           parentEventID         2444             0
149                        parentNameUsage         2444             0
142                      parentNameUsageID         2444             0
36                                 pathway         2444             0
156                                 phylum         2444          2444
193                              phylumKey         2444          2444
100                  pointRadiusSpatialFit         2444             0
39                            preparations         2444           716
51                 previousIdentifications         2444            34
209                              projectId         2444             0
204                               protocol         2444          2444
212                  publishedByGbifRegion         2444          2444
5                                publisher         2444             0
179                      publishingCountry         2444          2444
22                            recordNumber         2444            11
23                              recordedBy         2444          2431
24                            recordedByID         2444            36
6                               references         2444          1706
208               relativeOrganismQuantity         2444             0
207                            repatriated         2444          2444
30                   reproductiveCondition         2444             0
7                             rightsHolder         2444          1618
72                          sampleSizeUnit         2444             0
71                         sampleSizeValue         2444             0
73                          samplingEffort         2444             0
70                        samplingProtocol         2444            15
147                         scientificName         2444          2444
140                       scientificNameID         2444             0
28                                     sex         2444           793
200                                species         2444          2444
199                             speciesKey         2444          2444
168                        specificEpithet         2444          2444
63                          startDayOfYear         2444          2438
84                           stateProvince         2444          2444
161                              subfamily         2444             0
166                               subgenus         2444             0
198                            subgenusKey         2444             0
163                               subtribe         2444             0
159                            superfamily         2444             0
146                         taxonConceptID         2444             0
139                                taxonID         2444          1586
190                               taxonKey         2444          2444
171                              taxonRank         2444          2444
177                           taxonRemarks         2444             5
175                        taxonomicStatus         2444          2444
162                                  tribe         2444             0
8                                     type         2444            91
132                             typeStatus         2444             0
203                           typifiedName         2444             0
101               verbatimCoordinateSystem         2444           665
91                           verbatimDepth         2444             0
89                       verbatimElevation         2444             0
68                       verbatimEventDate         2444          1729
130                 verbatimIdentification         2444             0
55                           verbatimLabel         2444             0
88                        verbatimLocality         2444          1727
102                            verbatimSRS         2444             0
202                 verbatimScientificName         2444          2444
172                      verbatimTaxonRank         2444             0
173                         vernacularName         2444           136
90                           verticalDatum         2444             0
33                                vitality         2444            26
80                               waterBody         2444             0
65                                    year         2444          2444
```

###### Attributes Returned for eBird Records¶

This count was made before filters were applied

```
No eBird Basic Dataset was queried.
```

#### Citations¶

###### eBird¶

```
No eBird Basic Dataset was queried
```

###### GBIF¶

```
Citations-- 
When using this dataset please use the following citation and pay attention to the rights documented in rights.txt:
Austin J, Viani K, Hammond F, Massa M, Sharp S (2024). Historic Wildlife Roadkill Reports from Vermont, USA (1971-2006). Vermont Center for Ecostudies. Occurrence dataset https://doi.org/10.15468/6a4xjj accessed via GBIF.org on 2025-01-23.
Motz G (2025). Vertebrate Zoology Division - Mammalogy, Yale Peabody Museum. Yale University Peabody Museum. Occurrence dataset https://doi.org/10.15468/4mm6uc accessed via GBIF.org on 2025-01-23.
Cook J (2024). MSB Mammal Collection (Arctos). Version 35.94. Museum of Southwestern Biology. Occurrence dataset https://doi.org/10.15468/oirgxw accessed via GBIF.org on 2025-01-23.
Harvard University M, Morris P J (2025). Museum of Comparative Zoology, Harvard University. Version 162.454. Museum of Comparative Zoology, Harvard University. Occurrence dataset https://doi.org/10.15468/p5rupv accessed via GBIF.org on 2025-01-23.
Bentley A, Krejsa D (2024). KUBI Mammalogy Collection. Version 26.82. University of Kansas Biodiversity Institute. Occurrence dataset https://doi.org/10.15468/a3woj7 accessed via GBIF.org on 2025-01-23.
Garretson A, Napoli M, Feldsine N, Long E, Huth P, Smiley D, Forester A, Pierce E, Smiley S, Thompson J (2022). Mohonk Preserve Historical Observational Biodiversity Data. Mohonk Preserve. Occurrence dataset https://doi.org/10.15468/tckm2a accessed via GBIF.org on 2025-01-23.
California Academy of Sciences: CAS Mammalogy (MAM) https://doi.org/10.15468/dhbozg accessed via GBIF.org on 2025-01-23.
Kuprewicz E (2020). UConn Mammals. Version 3.3. University of Connecticut. Occurrence dataset https://doi.org/10.15468/dbs8w7 accessed via GBIF.org on 2025-01-23.
UMMZ Mammals Data Group, LSA IT A (2025). University of Michigan Museum of Zoology, Division of Mammals. Version 8.81. University of Michigan Museum of Zoology. Occurrence dataset https://doi.org/10.15468/dx3rcj accessed via GBIF.org on 2025-01-23.
Galbreath K (2024). Northern Michigan University (NMU) Mammal Specimens (Arctos). Version 1.84. Northern Michigan University. Occurrence dataset https://doi.org/10.15468/zazfgp accessed via GBIF.org on 2025-01-23.
Whitehouse R, Gerhard D (2021). NYSM Mammals. Version 16.2. New York State Museum (NYSM). Occurrence dataset https://doi.org/10.15468/awwifu accessed via GBIF.org on 2025-01-23.
Conroy C (2024). MVZ Mammal Collection (Arctos). Version 35.93. Museum of Vertebrate Zoology. Occurrence dataset https://doi.org/10.15468/uwudf9 accessed via GBIF.org on 2025-01-23.
iNaturalist contributors, iNaturalist (2025). iNaturalist Research-grade Observations. iNaturalist.org. Occurrence dataset https://doi.org/10.15468/ab3s5x accessed via GBIF.org on 2025-01-23.
University of Wisconsin – Stevens Point (2025). University of Wisconsin Stevens Point Mammals. Occurrence dataset https://doi.org/10.15468/7mp2c4 accessed via GBIF.org on 2025-01-23.
Bradley J (2025). UWBM Mammalogy Collection (Arctos). University of Washington Burke Museum. Occurrence dataset https://doi.org/10.15468/qziy3w accessed via GBIF.org on 2025-01-23.
```

```
Rights-- 
Dataset: Historic Wildlife Roadkill Reports from Vermont, USA (1971-2006) 
Rights as supplied: http://creativecommons.org/publicdomain/zero/1.0/legalcode
Dataset: Vertebrate Zoology Division - Mammalogy, Yale Peabody Museum 
Rights as supplied: http://creativecommons.org/publicdomain/zero/1.0/legalcode
Dataset: MSB Mammal Collection (Arctos) 
Rights as supplied: http://creativecommons.org/publicdomain/zero/1.0/legalcode
Dataset: Museum of Comparative Zoology, Harvard University 
Rights as supplied: http://creativecommons.org/licenses/by-nc/4.0/legalcode
Dataset: KUBI Mammalogy Collection 
Rights as supplied: http://creativecommons.org/licenses/by/4.0/legalcode
Dataset: Mohonk Preserve Historical Observational Biodiversity Data 
Rights as supplied: http://creativecommons.org/publicdomain/zero/1.0/legalcode
Dataset: CAS Mammalogy (MAM) 
Rights as supplied: http://creativecommons.org/publicdomain/zero/1.0/legalcode
Dataset: UConn Mammals 
Rights as supplied: http://creativecommons.org/publicdomain/zero/1.0/legalcode
Dataset: University of Michigan Museum of Zoology, Division of Mammals 
Rights as supplied: http://creativecommons.org/licenses/by-nc/4.0/legalcode
Dataset: Northern Michigan University (NMU) Mammal Specimens (Arctos) 
Rights as supplied: http://creativecommons.org/publicdomain/zero/1.0/legalcode
Dataset: NYSM Mammals 
Rights as supplied: http://creativecommons.org/publicdomain/zero/1.0/legalcode
Dataset: MVZ Mammal Collection (Arctos) 
Rights as supplied: http://creativecommons.org/publicdomain/zero/1.0/legalcode
Dataset: iNaturalist Research-grade Observations 
Rights as supplied: http://creativecommons.org/licenses/by-nc/4.0/legalcode
Dataset: University of Wisconsin Stevens Point Mammals 
Rights as supplied: http://creativecommons.org/licenses/by/4.0/legalcode
Dataset: UWBM Mammalogy Collection (Arctos) 
Rights as supplied: http://creativecommons.org/publicdomain/zero/1.0/legalcode
```

```
DOI-- 
https://doi.org/10.15468/dl.nkstk9
```

```
GBIF download key-- 
0003098-250121130708018
```

#### Runtime¶

```
0:01:31.079509
```

#### Optional Output¶

Remove "#" to activate desired statements.

```
[NbConvertApp] Converting notebook mFISHx0_Observations.ipynb to html
[NbConvertApp] Writing 970082 bytes to C:\Workspaces\RangeMaps\fisher\mFISHx0_Observations.html
```
