## Supplementary material for "A Framework for Transparent and Repeatable Species Range Maps": Occurrence Record Queries: File S2.html

mFISHx1\_Observations


### Fisher Records from GBIF¶

```
Notebook run 2025-01-23 13:45:03.423585
Results were saved in C:/Workspaces/RangeMaps/fisher/mFISHx1_Observations.sqlite
```

#### Taxon Concept¶

```
{'EBIRD_ID': 'None',
 'GBIF_ID': 5218855,
 'ID': 'mfishx1',
 'TAXON_EOO': None,
 'common_name': 'fisher',
 'detection_distance_m': 50,
 'scientific_name': 'Martes pennanti'}
```

#### Filter Set¶

```
Filter set name: Fisher1
```

###### GBIF Request Method¶

```
Request a Darwin Core Archive? True
```

###### Date Limit¶

Notes: The taxa concept was abandoned by the ASM taxonomy between 2012 and 2014.

```
Years: 2000,2015
Months: 1,12
```

###### Country¶

Notes:

```
Country: US
```

###### Bounding Box¶

Notes:

```
Latitude range: None
Longitude range: None
```

###### Area of Interest¶

Notes:

```
None
```

###### Taxon EOO¶

Notes:

```
Use taxon extent of occurrence? False
```

###### Geoissue¶

Notes:

```
Records with geoissues OK? False
```

###### Collections¶

Notes:

```
Omit: None
```

###### Institutions¶

Notes:

```
Omit: None
```

###### Datasets¶

Notes:

```
Omit: None
```

###### Coordinate Uncertainty¶

Notes:

```
Coordinate uncertainty required? False
Default coordinate uncertainty to use: None
Maximum allowable coordinate uncertainty: 100000
```

###### Bases¶

Notes:

```
Omit: None
```

###### Sampling Protocols¶

Notes:

```
Omit: None
```

###### Issues¶

Notes: Invalid dates could create errors in the removal of duplicates and/or assigning records to time periods. Taxon matching needs to be correct so that other species' records are not included. Invalid type statuses could allow the inclusion of fossils.

```
Omit: ['TAXON_MATCH_HIGHERRANK', 'TYPE_STATUS_INVALID', 'RECORDED_DATE_INVALID']
```

###### Duplicates¶

Notes:

```
Allow duplicates? False
```

###### Filter Set Summary¶

```
{'bases_omit': None,
 'collection_codes_omit': None,
 'country': 'US',
 'datasets_omit': None,
 'default_coordUncertainty': None,
 'duplicate_coord_date_OK': False,
 'geoissue': False,
 'get_dwca': True,
 'has_coordinate_uncertainty': False,
 'institutions_omit': None,
 'issues_omit': ['TAXON_MATCH_HIGHERRANK',
                 'TYPE_STATUS_INVALID',
                 'RECORDED_DATE_INVALID'],
 'lat_range': None,
 'lon_range': None,
 'max_coordinate_uncertainty': 100000,
 'months_range': '1,12',
 'name': 'Fisher1',
 'query_polygon': None,
 'sampling_protocols_omit': None,
 'use_taxon_geometry': False,
 'years_range': '2000,2015'}
```

#### Processing¶

```
Prepared filter set and sorted out geometry constraints: 0:00:00
175 records available
```

```
INFO:Your download key is 0003107-250121130708018
```

```
Waiting for the Darwin Core Archive.....
```

```
INFO:Download file size: 63026 bytes
INFO:On disk at C:/Workspaces/RangeMaps/fisher//0003107-250121130708018.zip
```

```
Wait time for DWcA creation: 0:01:15.078716
Wait time for DWcA download: 0:00:00.984526
Wait time for reading the DwCA: 0:00:00.238279
Stored GBIF Download DOI etc.: 0:00:00.005106
Summarized fields returned: 0:00:00.013999
Prepared GBIF records for processing: 0:00:00.033705
```

```
Prepared data frames for processing: 0:00:00.012100
Summarized values acquired: 0:00:00.064219
An unsupported date format has been found:  2001-01
An unsupported date format has been found:  2001-12
An unsupported date format has been found:  2005-10
An unsupported date format has been found:  2001-02
An unsupported date format has been found:  2001-02
Parsed event dates: 0:00:00.003000
Calculated nominal coordinate precision: 0:00:00.007552
Number of georeferenced GBIF records: 50
Approximating coordinate uncertanties for GBIF records
Prepared records and calculated radii:0:00:00.018557
Performed filtering: 0:00:00.007000
20 duplicate records dropped: 0:00:00.033303
Saved summary of filtering results: 0:00:00.096563
```

```
148 records were saved in the output database
```

#### Results of the Filtering¶

###### Sources¶

```
  institutionID collectionCode datasetName  acquired  removed  retained
0          UWBM           Mamm         nan        12        1        11
1           nan        Mammals         nan         2        1         1
2           nan             ZM         nan        54       17        37
3           nan            nan         nan       106        7        99
```

###### Bases¶

```
                value  acquired  removed  retained
0   HUMAN_OBSERVATION       106        7        99
1  PRESERVED_SPECIMEN        69       20        49
```

###### Protocols¶

```
  value  acquired  removed  retained
0   nan       175       27       148
```

###### Issues¶

```
                                                                                       value  acquired  removed  retained
0          CONTINENT_DERIVED_FROM_COORDINATES;INSTITUTION_MATCH_FUZZY;COLLECTION_MATCH_FUZZY        10        0        10
1  CONTINENT_DERIVED_FROM_COORDINATES;TAXON_MATCH_TAXON_ID_IGNORED;INSTITUTION_MATCH_FUZZ...         2        1         1
2  COORDINATE_REPROJECTED;CONTINENT_DERIVED_FROM_COORDINATES;INSTITUTION_MATCH_FUZZY;COLL...         2        1         1
3                 COORDINATE_ROUNDED;INSTITUTION_MATCH_FUZZY;INSTITUTION_COLLECTION_MISMATCH         3        0         3
4                                    INSTITUTION_MATCH_FUZZY;INSTITUTION_COLLECTION_MISMATCH        51       17        34
5                                           OCCURRENCE_STATUS_INFERRED_FROM_INDIVIDUAL_COUNT       106        7        99
```

Unique issues that were associated with records:

```
OCCURRENCE_STATUS_INFERRED_FROM_INDIVIDUAL_COUNT
COLLECTION_MATCH_FUZZY
COORDINATE_REPROJECTED
CONTINENT_DERIVED_FROM_COORDINATES
INSTITUTION_COLLECTION_MISMATCH
TAXON_MATCH_TAXON_ID_IGNORED
COORDINATE_ROUNDED
INSTITUTION_MATCH_FUZZY
```

###### Establishment Means¶

```
  value  acquired  removed  retained
0   nan       175       27       148
```

###### Identification Qualifiers¶

```
No identification qualifiers were reported.
```

#### Descriptions of Retained Records¶

###### Locations¶

###### Years Represented¶

Out[34]:

```
Text(0.5, 1.0, 'Occurrences per Year')
```

###### Months Represented¶

Out[35]:

```
Text(0.5, 1.0, 'Occurrences per Month')
```

###### Distribution of Coordinate Uncertainty Values for Retained Records¶

###### Distribution of Point-radius Values for Retained Records¶

Out[38]:

```
Text(0.5, 1.0, 'Compiled Point-radius Buffer Lengths')
```

###### Distribution of Nominal Coordinate Precisions for Retained Records¶

Out[39]:

```
Text(0.5, 1.0, 'Nominal Precisions of Coordinates')
```

###### Remarks¶

```
General remarks:
<NA>
```

```
Event remarks:
<NA>
native; missing all feet; neck circ=180mm
native; "Phillips kit male"; ear punch dry & refilled JAN05
native; "Phillip's Cr. F - 21"
native; "Adult Male M-24"
native; "Curr Creek F-22"
native; "Quarry F-20"; ear punch dry & refilled JAN 2005
native; "Rustler Peak F-18"
native; "New Guy M-23"
native; "Middlefork F-19"; ear punch dry & refilled JAN05
native; "Foster Cr male - 02"; flat skin
native; "Union Creek Female-11"; necropsied by collector
```

```
Occurrence remarks:
<NA>
remote area but some houses // Skin off when measurements taken; stomach contents: Vitis labrusca 10%, Sylviagus floridanus 90%
Swampy area // Skin off when measurements taken; stomach contents: Scalopsis aquaticus 100%.
Woodland // Skin off when measurements taken; stomach contents: 74+ Dolichovespula germanica, tail and two feet from a Peromyscus sp., tail from a Tamias striatus.
caught in culvert // Skin off when measurements taken.
Woodland // Skin off when measurements taken.
Roadkill; measurements: 855-336-120-45-2350g
```

```
Location remarks:
<NA>
from collector using Terrain Navigator Pro; VERBATIMELEVATION: 1268; GEOREFERENCEVERIFICATIONSTATUS: requires verification
Converted from collector's UTM by Graphical Locator; NAD27; VERBATIMELEVATION: 1036; GEOREFERENCEVERIFICATIONSTATUS: requires verification
From collector's TRS by Graphical Locator; VERBATIMELEVATION: 1034; VERBATIMLATITUDE: 423682N; VERBATIMLONGITUDE: 1222711W; GEOREFERENCEVERIFICATIONSTATUS: requires verification
Converted from collector's UTM by Graphical Locator; NAD27; VERBATIMELEVATION: 1705; GEOREFERENCEVERIFICATIONSTATUS: requires verification
Converted from collector's UTM by Graphical Locator; NAD27; VERBATIMELEVATION: 1342; GEOREFERENCEVERIFICATIONSTATUS: requires verification
Converted from collector's UTM by Graphical Locator; NAD27; VERBATIMELEVATION: 1287; GEOREFERENCEVERIFICATIONSTATUS: requires verification
Converted from collector's UTM by Graphical Locator; NAD27; VERBATIMELEVATION: 1266; GEOREFERENCEVERIFICATIONSTATUS: requires verification
Converted from collector's UTM by Graphical Locator; NAD27; VERBATIMELEVATION: 1209; GEOREFERENCEVERIFICATIONSTATUS: requires verification
Converted from collector's UTM by Graphical Locator; NAD27; VERBATIMELEVATION: 793; GEOREFERENCEVERIFICATIONSTATUS: requires verification
Converted from collector's TRS by Graphical Locator Homepage; VERBATIMELEVATION: 1034; VERBATIMLATITUDE: 424726N; VERBATIMLONGITUDE: 1223418W; GEOREFERENCEVERIFICATIONSTATUS: requires verification
Converted from Collector's TRS by Graphical Locator Homepage; VERBATIMELEVATION: 1097; GEOREFERENCEVERIFICATIONSTATUS: requires verification
```

```
Identified remarks:
<NA>
```

```
Georeference remarks:
<NA>
```

###### Attributes Returned for GBIF Records¶

This count was made before filters were applied

```
                                 attribute  included(n)  populated(n)
Field                                                                
148                      acceptedNameUsage          175             0
141                    acceptedNameUsageID          175             0
201                 acceptedScientificName          175           175
191                       acceptedTaxonKey          175           175
0                             accessRights          175            67
41                   associatedOccurrences          175             0
50                     associatedOrganisms          175             0
42                    associatedReferences          175             0
43                     associatedSequences          175             0
44                          associatedTaxa          175             0
16                           basisOfRecord          175           175
128                                    bed          175             0
32                                behavior          175            70
1                    bibliographicCitation          175            54
31                                   caste          175             0
21                           catalogNumber          175            69
157                                  class          175           175
194                               classKey          175           175
13                          collectionCode          175            69
10                            collectionID          175            15
79                               continent          175           175
99                     coordinatePrecision          175             0
98           coordinateUncertaintyInMeters          175            55
83                             countryCode          175           175
85                                  county          175            66
170                        cultivarEpithet          175             0
18                     dataGeneralizations          175           106
11                               datasetID          175             0
178                             datasetKey          175           175
14                             datasetName          175             0
135                         dateIdentified          175            13
67                                     day          175           170
96                         decimalLatitude          175           175
97                        decimalLongitude          175           175
35                   degreeOfEstablishment          175             0
183                                  depth          175             0
184                          depthAccuracy          175             0
40                             disposition          175             0
185           distanceFromCentroidInMeters          175             0
19                       dynamicProperties          175            13
120               earliestAgeOrLowestStage          175             0
112            earliestEonOrLowestEonothem          175             0
118            earliestEpochOrLowestSeries          175             0
114             earliestEraOrLowestErathem          175             0
116           earliestPeriodOrLowestSystem          175             0
181                              elevation          175            17
182                      elevationAccuracy          175            17
64                            endDayOfYear          175           170
34                      establishmentMeans          175             0
61                               eventDate          175           175
57                                 eventID          175             0
75                            eventRemarks          175            12
62                               eventTime          175            13
59                               eventType          175             0
160                                 family          175           175
196                              familyKey          175           175
74                              fieldNotes          175             0
60                             fieldNumber          175             0
104                           footprintSRS          175             0
105                    footprintSpatialFit          175             0
103                           footprintWKT          175            13
126                              formation          175             0
211                             gbifRegion          175           175
165                            genericName          175           175
164                                  genus          175           175
197                               genusKey          175           175
222                          geodeticDatum          175           175
111                    geologicalContextID          175             0
108                   georeferenceProtocol          175            67
110                    georeferenceRemarks          175             0
109                    georeferenceSources          175            15
37          georeferenceVerificationStatus          175            54
106                        georeferencedBy          175            13
107                      georeferencedDate          175            13
125                                  group          175             0
69                                 habitat          175             0
188                          hasCoordinate          175           175
189                    hasGeospatialIssues          175           175
154                   higherClassification          175            69
78                         higherGeography          175            67
77                       higherGeographyID          175             0
123            highestBiostratigraphicZone          175             0
129                       identificationID          175             0
131                identificationQualifier          175            13
136               identificationReferences          175             0
138                  identificationRemarks          175            13
137       identificationVerificationStatus          175             0
133                           identifiedBy          175            13
134                         identifiedByID          175             0
25                         individualCount          175           160
17                     informationWithheld          175             0
167                    infragenericEpithet          175             0
169                   infraspecificEpithet          175             0
12                         institutionCode          175            69
9                            institutionID          175            13
210                            isSequenced          175           175
82                                  island          175             0
81                             islandGroup          175             0
186                                  issue          175           175
221                    iucnRedListCategory          175           175
155                                kingdom          175           175
192                             kingdomKey          175           175
2                                 language          175            67
206                            lastCrawled          175           175
180                        lastInterpreted          175           175
205                             lastParsed          175           175
121                latestAgeOrHighestStage          175             0
113             latestEonOrHighestEonothem          175             0
119             latestEpochOrHighestSeries          175             0
115              latestEraOrHighestErathem          175             0
117            latestPeriodOrHighestSystem          175             0
213                              level0Gid          175           175
214                             level0Name          175           175
215                              level1Gid          175           175
216                             level1Name          175           175
217                              level2Gid          175           175
218                             level2Name          175           175
219                              level3Gid          175             0
220                             level3Name          175             0
3                                  license          175           175
29                               lifeStage          175           107
124                lithostratigraphicTerms          175             0
87                                locality          175           175
94                     locationAccordingTo          175            13
76                              locationID          175             0
95                         locationRemarks          175            13
122             lowestBiostratigraphicZone          175             0
53                        materialEntityID          175             0
54                   materialEntityRemarks          175             0
56                        materialSampleID          175             0
93     maximumDistanceAboveSurfaceInMeters          175             0
187                              mediaType          175             2
127                                 member          175             0
92     minimumDistanceAboveSurfaceInMeters          175             0
4                                 modified          175            69
66                                   month          175           175
86                            municipality          175             0
151                        nameAccordingTo          175             0
144                      nameAccordingToID          175             0
152                        namePublishedIn          175             0
145                      namePublishedInID          175             0
153                    namePublishedInYear          175             0
174                      nomenclaturalCode          175            67
176                    nomenclaturalStatus          175             0
20                            occurrenceID          175           175
46                       occurrenceRemarks          175             6
38                        occurrenceStatus          175           175
158                                  order          175           175
195                               orderKey          175           175
47                              organismID          175            13
48                            organismName          175             0
26                        organismQuantity          175             0
27                    organismQuantityType          175             0
52                         organismRemarks          175             0
49                           organismScope          175             0
150                      originalNameUsage          175             0
143                    originalNameUsageID          175             0
45                     otherCatalogNumbers          175            12
15                    ownerInstitutionCode          175             0
58                           parentEventID          175             0
149                        parentNameUsage          175             0
142                      parentNameUsageID          175             0
36                                 pathway          175             0
156                                 phylum          175           175
193                              phylumKey          175           175
100                  pointRadiusSpatialFit          175             0
39                            preparations          175            58
51                 previousIdentifications          175            13
209                              projectId          175             0
204                               protocol          175           175
212                  publishedByGbifRegion          175           175
5                                publisher          175             0
179                      publishingCountry          175           175
22                            recordNumber          175             2
23                              recordedBy          175           163
24                            recordedByID          175             0
6                               references          175            69
208               relativeOrganismQuantity          175             0
207                            repatriated          175           175
30                   reproductiveCondition          175             0
7                             rightsHolder          175             0
72                          sampleSizeUnit          175             0
71                         sampleSizeValue          175             0
73                          samplingEffort          175             0
70                        samplingProtocol          175             0
147                         scientificName          175           175
140                       scientificNameID          175             0
28                                     sex          175            71
200                                species          175           175
199                             speciesKey          175           175
168                        specificEpithet          175           175
63                          startDayOfYear          175           170
84                           stateProvince          175           175
161                              subfamily          175             0
166                               subgenus          175             0
198                            subgenusKey          175             0
163                               subtribe          175             0
159                            superfamily          175             0
146                         taxonConceptID          175             0
139                                taxonID          175             2
190                               taxonKey          175           175
171                              taxonRank          175           175
177                           taxonRemarks          175             1
175                        taxonomicStatus          175           175
162                                  tribe          175             0
8                                     type          175            67
132                             typeStatus          175             0
203                           typifiedName          175             0
101               verbatimCoordinateSystem          175            13
91                           verbatimDepth          175             0
89                       verbatimElevation          175             0
68                       verbatimEventDate          175            69
130                 verbatimIdentification          175             0
55                           verbatimLabel          175             0
88                        verbatimLocality          175            67
102                            verbatimSRS          175             0
202                 verbatimScientificName          175           175
172                      verbatimTaxonRank          175             0
173                         vernacularName          175           106
90                           verticalDatum          175             0
33                                vitality          175             0
80                               waterBody          175             0
65                                    year          175           175
```

###### Attributes Returned for eBird Records¶

This count was made before filters were applied

```
No eBird Basic Dataset was queried.
```

#### Citations¶

###### eBird¶

```
No eBird Basic Dataset was queried
```

###### GBIF¶

```
Citations-- 
When using this dataset please use the following citation and pay attention to the rights documented in rights.txt:
Whitehouse R, Gerhard D (2021). NYSM Mammals. Version 16.2. New York State Museum (NYSM). Occurrence dataset https://doi.org/10.15468/awwifu accessed via GBIF.org on 2025-01-23.
Garretson A, Napoli M, Feldsine N, Long E, Huth P, Smiley D, Forester A, Pierce E, Smiley S, Thompson J (2022). Mohonk Preserve Historical Observational Biodiversity Data. Mohonk Preserve. Occurrence dataset https://doi.org/10.15468/tckm2a accessed via GBIF.org on 2025-01-23.
University of Wisconsin – Stevens Point (2025). University of Wisconsin Stevens Point Mammals. Occurrence dataset https://doi.org/10.15468/7mp2c4 accessed via GBIF.org on 2025-01-23.
Bradley J (2025). UWBM Mammalogy Collection (Arctos). University of Washington Burke Museum. Occurrence dataset https://doi.org/10.15468/qziy3w accessed via GBIF.org on 2025-01-23.
Galbreath K (2024). Northern Michigan University (NMU) Mammal Specimens (Arctos). Version 1.84. Northern Michigan University. Occurrence dataset https://doi.org/10.15468/zazfgp accessed via GBIF.org on 2025-01-23.
```

```
Rights-- 
Dataset: NYSM Mammals 
Rights as supplied: http://creativecommons.org/publicdomain/zero/1.0/legalcode
Dataset: Mohonk Preserve Historical Observational Biodiversity Data 
Rights as supplied: http://creativecommons.org/publicdomain/zero/1.0/legalcode
Dataset: University of Wisconsin Stevens Point Mammals 
Rights as supplied: http://creativecommons.org/licenses/by/4.0/legalcode
Dataset: UWBM Mammalogy Collection (Arctos) 
Rights as supplied: http://creativecommons.org/publicdomain/zero/1.0/legalcode
Dataset: Northern Michigan University (NMU) Mammal Specimens (Arctos) 
Rights as supplied: http://creativecommons.org/publicdomain/zero/1.0/legalcode
```

```
DOI-- 
https://doi.org/10.15468/dl.hsuv5g
```

```
GBIF download key-- 
0003107-250121130708018
```

#### Runtime¶

```
0:01:23.142036
```

#### Optional Output¶

Remove "#" to activate desired statements.

```
[NbConvertApp] Converting notebook mFISHx1_Observations.ipynb to html
[NbConvertApp] Writing 871018 bytes to C:\Workspaces\RangeMaps\fisher\mFISHx1_Observations.html
```
